# TLL1 knockdown attenuates prostate cancer progression by enhancing antitumor immunity

**DOI:** 10.1101/2024.11.11.623110

**Authors:** Jing-Lan Hao, Jia-Qi He, Hang Hu, Xiao-Yu Wu, Zi-Heng Zhu, Xi Zhao, Lan Li, Yong-Tong Ruan, Juan Yang, Ming Fu, Kai Zhang, Ping Gao, Xiao-Ming Dong

## Abstract

Due to the inevitable progression to castration-resistant prostate cancer (CRPC) following treatment with androgen deprivation therapy (ADT), it is essential to develop novel treatment approaches for managing CRPC. Here we showed that metalloproteinase TLL1 was positively associated with prostate cancer aggressiveness. Mechanistically, TLL1 promoted prostate cancer cells migration and metastasis through cleaving latent TGF-β1 to activate TGF-β signaling pathway. Moreover, *LINC01179* interacted with Miz1 to attenuate TLL1 expression and *LINC01179* impaired prostate cancer cell proliferation and migration ability by suppressing TLL1 expression to deactivate TGF-β signaling activity. Meanwhile, we observed that TLL1 increased the expression of PD-L1 by activating TGF-β signaling pathway and TLL1 depletion enhanced the antitumor efficacy by anti-PD-1 antibody via augmenting the infiltration proportions of CD8^+^ T cells in tumors. In addition, T cell-specific overexpression of TLL1 disrupted T cell development in the thymus. TLL1 overexpression in T cells accelerated RM-1 prostate tumor growth in mice by decreasing the infiltration of CD8^+^ T cells into tumors. Collectively, our results revealed that TLL1 may be a potential therapeutic target to alter prostate cancer progression.

## INTRODUCTION

Prostate cancer (PCa) is the leading malignant tumor in males and it will progress into the lethal CRPC state within a few months to 2-3 years after ADT [1]. During the last decades, several crucial factors including transforming growth factor-β (TGF-β) have been found to drive the emergence of CRPC [2–4]. Previous studies have reported that TGF-β signaling can be activated during the ADT process in PCa [5,6]. To date, the mechanisms of how TGF-β signaling was activated during prostate cancer initiation and progression are still elusive.

Programmed death-ligand 1 (PD-L1), encoded by CD274 gene, is a transmembrane protein that facilitates immune evasion by interacting with the inhibitory receptor programmed cell death protein 1 (PD-1) on T cells, leading to T-cell functional exhaustion [7,8]. A variety of cancer cells, including those from melanoma, lung, bladder, breast, and prostate cancers, increase the expression of PD-L1 as a mechanism to escape immune surveillance. Strategies that target immune checkpoints, like the interaction between PD-L1 and its receptor PD-1, have received approval for the treatment of various human cancers, offering significant clinical advantages [9]. However, many cancer patients, especially prostate cancer, fail to respond to the immune treatment with anti-PD-1 or anti-PD-L1 antibodies [10] *(10)*. Recent advances in cancer immune therapies revealed that response to anti-PD-1/PD-L1 treatment might correlate with the PD-L1 expression levels in cancer cells [11–13]. Therefore, it is crucial to explore the pathways controlling PD-L1 protein expression, which is critical for formulating treatment approaches to enhance the effectiveness of immunotherapy for prostate cancer.

TLL1 is a zinc ion dependent metalloproteinase, highly homologous to bone morphogenetic protein 1 (BMP1) and located in the extracellular matrix [14,15]. Accumulated evidence has shown a close correlation between TGF-β/BMP signaling pathway and bone metastasis in prostate cancer [16,17]. Similar in function to BMPs, studies have reported that TLL1 can affect the differentiation of liver stem cells by regulating the activity of TGF-β signaling pathways [18]. Besides, TLL1 could modify the tumor microenvironment by cleaving ANGPTL2 into fragments to inhibit osteosarcoma cell migration and metastasis [19]. However, the role of TLL1 in prostate cancer has not yet been studied.

Long-non coding RNA (LncRNA) has been found to play important roles in numerous biological processes [20]. Many studies proved that long-non coding RNAs participated in tumor initiation and progression, act as either oncogenes or tumor suppressors [21,22]. Some of them could interact with transcription factors to disturb their binding affinity to other proteins or DNA fragments [23]. It was reported that Myc-interacting zinc finger protein 1 (Miz1) together with Myc mediated transrepressional activities. Myc-Miz1 and TGF-β signaling antagonistically regulated the expression of a cell cycle inhibitory protein p15 [24–26]. Until now, whether there have any long-non coding RNAs involved in this antagonistic regulating relationship between Miz1 and TGF-β signaling is still unclear.

In the present study, we showed that TLL1 was highly expressed in prostate tumor samples and associated with shorter survival of patients. TLL1 enhanced prostate cancer cell migration and metastasis ability through cleaving latent TGF-β1 to activate TGF-β signaling pathway. Further, the long-non coding RNA *LINC01179* suppressed TLL1 expression by promoting Miz1 binding at the *TLL1* promoter region. *LINC01179* suppressed the proliferation and migration ability of prostate cancer cells by inhibiting TLL1 expression to deactivate TGF-β signaling pathway. Moreover, TLL1 increased the abundance of PD-L1 by activating TGF-β signaling pathway and TLL1 depletion enhanced the antitumor efficacy via augmenting the infiltration proportions of CD8^+^ T cells in tumors. In addition, TLL1 overexpression in T cells impaired T cell development in the thymus. TLL1 overexpression in T cells accelerated RM-1 prostate tumor growth in mice by decreasing the infiltration of CD8^+^ T cells into tumors. Taken together, we systemically analyzed the functional role of TLL1 in prostate cancer and suggested that TLL1 may be a potential therapeutic target for cancer immunotherapy.

## RESULTS

### TLL1 plays an oncogenic role in prostate cancer

To investigate the expression of TLL1 in prostate cancer, we first collected 77 pairs of prostate cancer tissues and the adjacent normal tissues, immunohistochemical (IHC) staining results showed that TLL1 was overexpressed in prostate cancer tissues compared with normal tissues (Fig. 1, A and B). Moreover, the protein levels of TLL1 were highly expressed in high Gleason score 9-10 compared with low Gleason score 6-8 prostate cancer tissues (Fig. 1C) and the elevated TLL1 expression was associated with shorter survival of patients (Fig. 1D). To further explore the *TLL1* mRNA level in prostate cancer, we analyzed the published prostate cancer cohort datasets and found that the *TLL1* mRNA level was up-regulated in high Gleason score 5 prostate cancer tissues compared with low Gleason score 3-4 tissues (Fig. 1E) and associated with shorter overall survival and disease-free survival of patients (Fig. 1F and fig. S1A). These results consist with the findings what we have found in the 77 pairs of prostate cancer tissues and suggested that TLL1 was associated with prostate cancer progression.

**Fig. 1.**
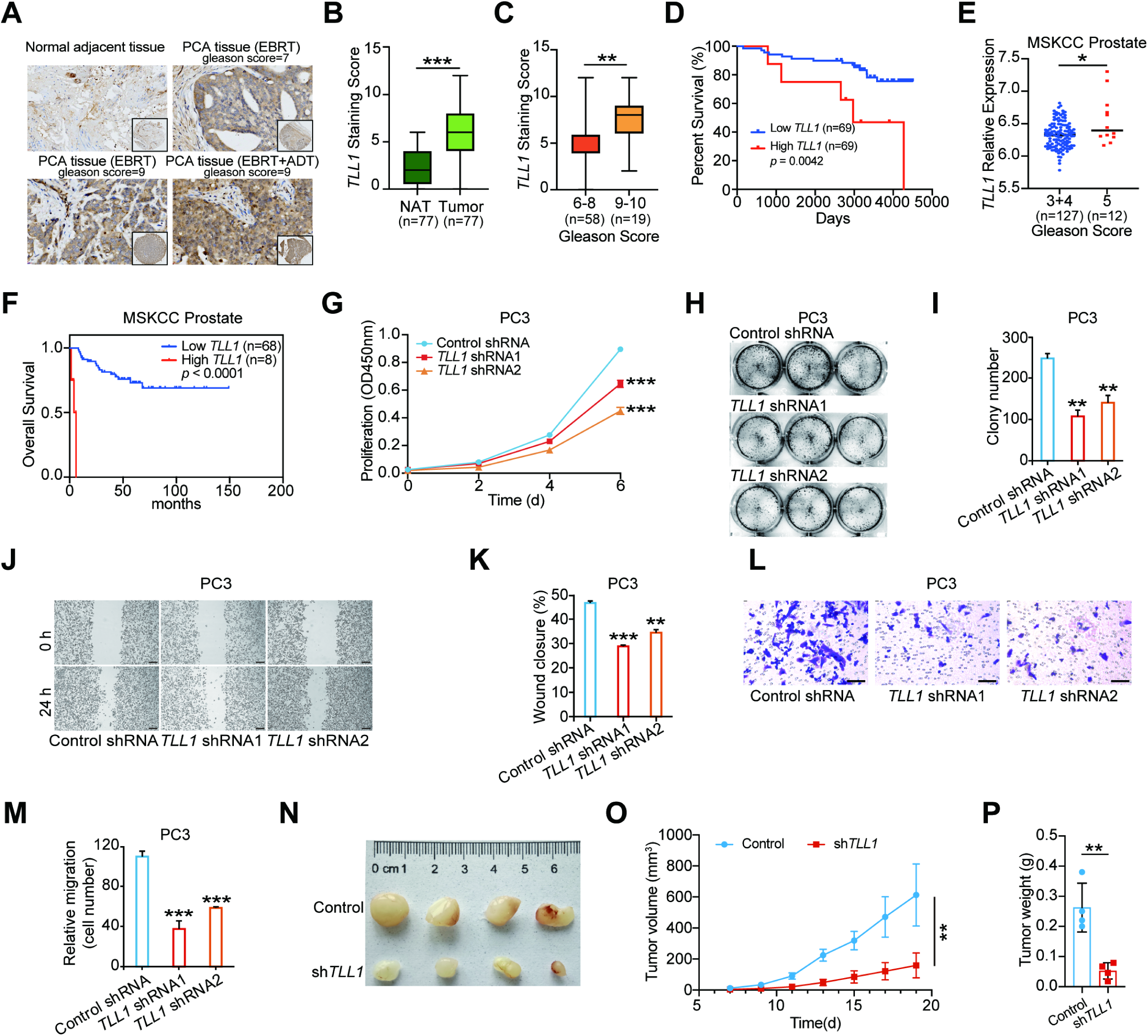
TLL1 plays an oncogenic role in prostate cancer. **(A)** Representative picture of TLL1 protein expression in prostate cancer tissue chip detected by IHC. **(B)** Quantification of TLL1 protein levels in prostate cancer tissue. (**C**) The correlation between TLL1 protein levels and Gleason score from prostate cancer tissue chip. (**D**) Association between overall survival of prostate cancer patients and TLL1 protein levels. (**E**) The correlation between *TLL1* expression and Gleason score from MSKCC Prostate data. (**F**) Association between overall survival of prostate cancer patients and *TLL1* expression levels from MSKCC Prostate database. (**G**) Effects of TLL1 depletion on cell proliferation measured by CCK8 assays in PC3 cells. (**H** and **I**) Representative pictures (H) and quantification analysis (I) of colony formation assays in control and TLL1 knockdown PC3 cells. (**J** and **K**) Representative pictures (J) and quantification analysis (K) of wound healing assays in control and TLL1 depleted PC3 cells. Scale bar = 200 μm. (**L** and **M**) Representative pictures (L) and quantification analysis (M) of migration assays in control or TLL1 knockdown PC3 cells. Scale bar = 100 μm. (**N** to **P**) Control and TLL1 knockdown PC3 cells were subcutaneously injected into BALB/c nude mice for xenograft growth (n = 4 per group). Tumor images (N), tumor volumes (O) and tumor weight (P) are shown. \**p* < 0.05, \*\**p* < 0.01, \*\*\**p* < 0.001. Data are shown as mean ± SD with three times of independent assay in **G**, **I**, **K** and **M**. In **B**, **C**, **E**, **G**, **I**, **K, M, O** and **P**, *p* values were assessed using two-tailed Student’s *t* tests. In **D** and **F**, *p* values were assessed using log rank test.

To further evaluate the functional role of TLL1 in prostate cancer progression, we established prostate cancer cell lines PC3 and 22Rv1 stably shRNA against *TLL1* (fig. S1, B and C), and the effect of TLL1 knockdown on cell proliferation was measured. CCK-8 assay was utilized to detect cell proliferation activity. The results showed that knockdown of TLL1 in PC3 and 22Rv1 cells significantly attenuated cell growth compared with control group (Fig. 1G and fig.S1D). Colony formation assay found that the colony formation ability of PC3 and 22Rv1 cells was inhibited after reducing the expression level of TLL1 (Fig. 1, H and I, and fig. S1, E and F). Further, wound healing assay showed that TLL1 suppression in PC3 cells restrained cell migration ability (Fig,1, J and K). The trans-well assay also indicated that inhibition of TLL1 expression in PC3 and 22Rv1 cells could also apparently weaken cell migration ability (Fig. 1, L and M, and fig. S1, G and H). Accordingly, TLL1 overexpression in PC3 facilitated cell migration ability (fig, S1, I to K).

Next, tumor xenograft models were established in nude mice to evaluate the effects of TLL1 on PC3 cells growth. As shown in Fig. 1N, the isolated tumors in the shTLL1 group were smaller than those in the control group. Furthermore, the tumor volume and weight were apparently decreased in the shTLL1 group compared to the control group (Fig. 1, O and P). Collectively, these results revealed that TLL1 was a novel oncogene and was positively associated with aggressive prostate cancer.

### TLL1 facilitated PCa cells migration and metastasis via cleaving latent TGF-β1 to enhance TGF-β signaling pathway

Considering TLL1 is the homologous gene of BMP1 and TLL1 could regulate TGF-β signaling pathway during hepatic differentiation process [27], we speculated that TLL1 might promote prostate cancer progression by activating TGF-β signaling pathway. We firstly analyzed the impact of TLL1 on the concentration of active TGF-β1, the results showed that TGF-β1 levels were decreased in TLL1 inhibited PCa cells and increased in TLL1 overexpressed PCa cells culture supernatant (Fig. 2, A and B, and fig. S2, A and B). It was known that BMP1 can cleave latent (TGF-β)-binding proteins. Following cleavage, the TGFβ precursor becomes active TGF-β [28]. We speculated that TLL1 as metalloproteinase might cleave latent TGF-β1 and activate TGF-β signaling pathway. To assess this point, we harvested TLL1 overexpression or knockdown PC3 cells culture supernatant and analyzed levels of TGF-β1 from concentrated supernatant, respectively. The results revealed that TLL1 overexpression increased the active TGF-β1 levels compared with control cells; TLL1 knockdown can lead to decreased levels of active TGF-β1 compared with control cells (Fig. 2,C and D). It was reported that activation of TGF-β signaling pathway could promote CRPC by inducing epithelial-mesenchymal transition (EMT) [3], thus we measured the protein levels of E-cadherin and Vimentin in TLL1 knockdown PCa cells. The results showed that reduced TLL1 expression apparently attenuated vimentin protein levels and enhanced E-cadherin expression levels (Fig. 2E). Moreover, we evaluated the impact of TLL1 on the phosphorylated Smad2 levels to determine whether TLL1 could activate TGF-β signaling pathway. The results indicated that TGF-β induced a significant increase of Smad2 phosphorylation, and TLL1 knockdown in PCa cells dramatically decreased the phosphorylated Smad2 levels compared with control cells (Fig. 2F). Consistently, forced expression of TLL1 in PCa cells strongly increased the phosphorylated Smad2 levels compared with the control group, which was blocked by TGF-β signaling inhibitor (LY-2109761) treatment (Fig. 2G, and fig. S2C).

**Fig. 2.**
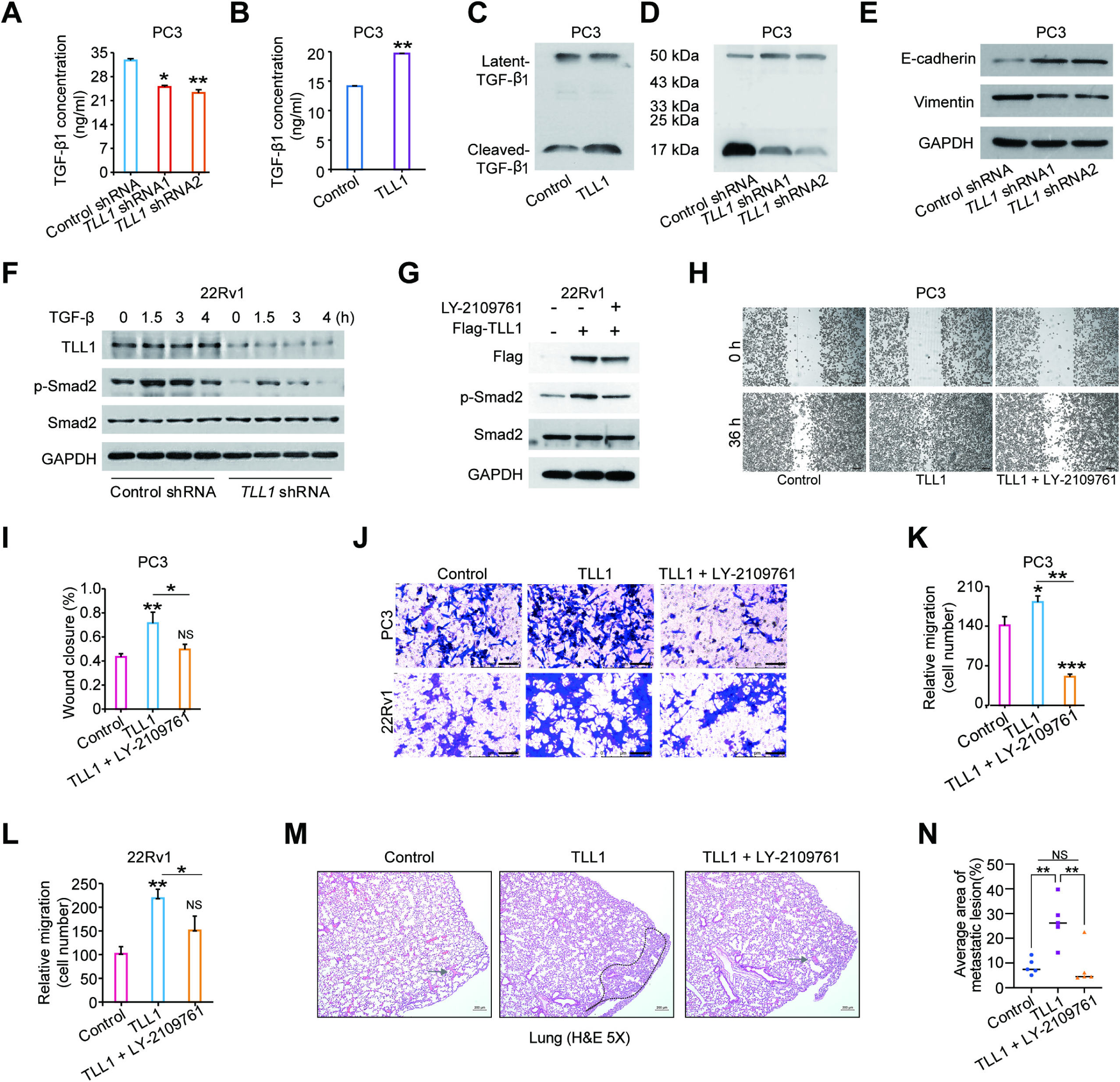
TLL1 facilitated PCa cells migration and metastasis via cleaving latent TGF-β1 to enhance TGF-β signaling pathway. **(A)** TGF-β1 concentration was detected by ELISA assay in control and TLL1 knockdown PC3 cells. **(B)** ELISA assay analysis of TGF-β1 concentration in control and TLL1 overexpressed PC3 cells. (**C**) The culture medium from TLL1 overexpressed PC3 cells and control cells was harvested and concentrated as described in materials and methods. Secreted latent TGF-β and activated TGF-β were visualized by western blotting. (**D**) The culture medium from TLL1 knockdown PC3 cells and control cells was harvested and concentrated as described in materials and methods. Secreted latent TGF-β and activated TGF-β were examined by western blotting. (**E**) Western blotting analysis of E-cadherin and Vimentin protein levels in control and TLL1 depleted PC3 cells. GAPDH was used as a loading control. (**F**) Western blot assay was performed to detect the effect of TLL1 knockdown on Smad2 phosphorylation level stimulated by TGF-β factors (10 µg/µL) in 22Rv1 cells. (**G**) TGF-β receptor inhibitor LY-2109761(2 µM) was added in TLL1 overexpressed 22Rv1 cells, the phosphorylated level of Smad2 were detected by western blot assay. (**H** and **I**) Representative pictures (H) and quantification analysis (I) of wound healing assays in control and TLL1 overexpressed PC3 cells treated with LY-2109761(2 μM). Scale bar = 200 μm. (**J** to **L**) Representative pictures (J) and quantification analysis (K-L) of migration assays in control and TLL1 overexpressed PCa cells stimulated with LY-2109761(2 μM). Scale bar = 100 μm. (**M** and **N**) Representative H&E images of lung metastases in all three groups of chemical castrated mice injection with LY-2109761 at 5× magnification (M). Scale bar = 200 μm. Scatter plot represents an average area in percentage of lung metastatic lesions in the aforementioned groups (N), n = 5 per group. NS, not significant, \**p* < 0.05, \*\**p* < 0.01, \*\*\**p* < 0.001. In **A**, **B**, **I**, **K and L**, data shown are mean ± SD of triplicate experiments. In **A**, **B**, **I**, **K**, **L** and **N**, *p* values were assessed using two-tailed Student’s *t* tests.

It has been well known that inhibition of TGF-β signaling could prevent the spreading of cancer cells [29]. The TGF-β signaling inhibitor is supposed to inverse the biological functions of TLL1 if it works by activating TGF-β signaling in PCa cells. Thus, TLL1 overexpressed PCa cells or control cells were treated with TGF-β pathway inhibitor LY-2109761 to evaluate whether LY-2109761 could inhibit cell migration caused by TLL1. The wound-healing and trans-well results indicated that the high migration ability of TLL1 overexpressed PCa cells was restrained by LY-2109761 treatment (Fig. 2, H to L, and fig. S2, D and E). Moreover, we used the inhibitor to treat the nude mouse which had been injected with TLL1 overexpressed PC3 cells or control cells, the H&E results showed that forced expression of TLL1 had more large-sized metastatic tumor lesions in the lung than the control group, which was impeded by LY-2109761 treatment (Fig. 2, M and N). Collectively, our results demonstrated that enhanced TGF-β signaling is a critical mediator of TLL1 overexpression–induced EMT and prostate cancer cell migration and metastasis, which can be effectively blocked by LY-2109761. Altogether, these results indicated that highly expressed TLL1 facilitates migration and metastasis of PCa cells by activating TGF-β signaling pathway.

### *LINC01179* interacted with Miz1 to regulate TLL1 expression

In order to reveal upstream regulatory molecules of TLL1 in prostate cancer, we observed a novel long-non coding RNA *LINC01179* on the genome, which is just located at the upstream of *TLL1*. To evaluate whether *LINC01179* can regulate TLL1 expression, we constructed stable *LINC01179* inhibition or overexpression cell strains in PCa. RT-qPCR and western blot assays were performed to detect TLL1 levels. The results showed that *LINC01179* inhibition led to the increased mRNA and protein levels of TLL1; *LINC01179* overexpression attenuated the mRNA and protein levels of TLL1 (Fig. 3, A to D). To determine the functional role of *LINC01179* in prostate cancer, we first confirmed that *LINC01179* is mainly located in the nucleus of PCa cells with a potential role in the regulation of gene expression (Fig. 3, E and F), then we performed native RNA pull-down assay using biotinylated DNA probes against *LINC01179*. Silver staining of the pull-down protein complexes identified two unique protein bands at ∼100 kDa compared with the control pull-down (Fig. 3G). Mass spectrometry analysis identified the lower band as Miz1 (fig. S3, A and B). We then validated this interaction by western blotting for Miz1 after *LINC01179* RNA pull-down. The results showed that there is a direct interaction between *LINC01179* and Miz1 (Fig. 3H). Further, we predicted the potential transcription factors binding at the TLL1 upstream and downstream regions using published ChIP-seq data (http://cistrome.org/). We observed that Miz1 might be the potential candidate transcription factor directly binding at *TLL1* (Fig. 3I). ChIP-qPCR assay further validated that Miz1 can strongly bind at the intron 1 of *TLL1* in the PCa cells and the binding affinity of Miz1 at the *TLL1* intron1 region was significantly halted after diminishing the expression of *LINC01179* (Fig. 3J). The luciferase assay showed that Miz1 inhibited the activity of *TLL1* promoter by binding to enhancer region of *TLL1* (Fig. 3K). Correlation analysis showed that *TLL1* levels were negatively correlated with *Miz1* expression (Fig. 3, L and M). Moreover, knockdown of Miz1 in PC3 cells increased the mRNA and protein levels of TLL1, accordingly, Miz1 overexpression decreased TLL1 expression (Fig. 3, N to P). Taken together, these results indicated that *LINC01179* interacted with Miz1 to suppress TLL1 expression levels.

**Fig. 3.**
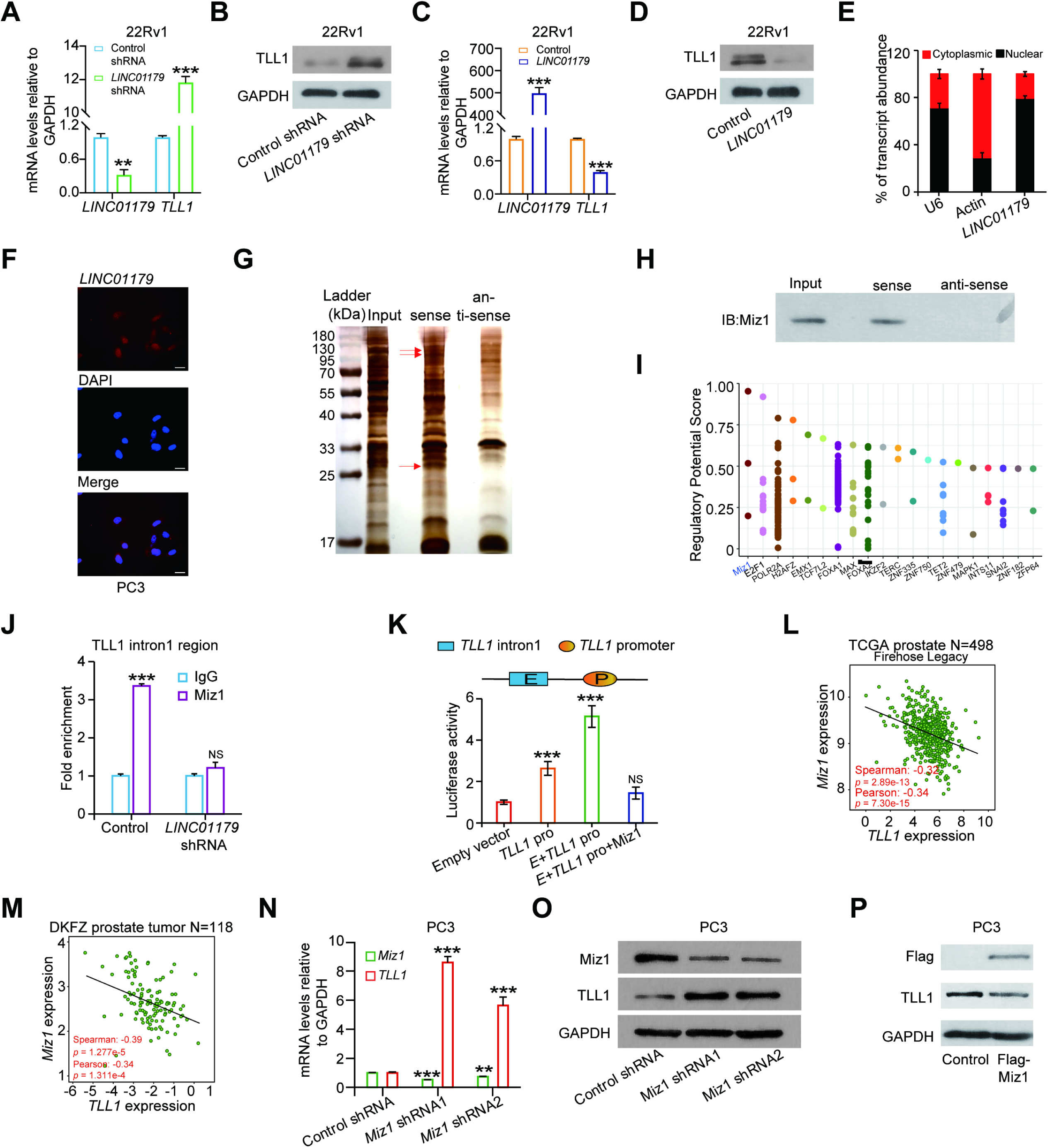
LINC01179 interacted with Miz1 to regulate TLL1 expression. (**A**) qRT-PCR analysis of the mRNA levels of *TLL1* in control and LINC01179 depleted 22Rv1 cells. (**B**) Western blotting analysis of TLL1 expression levels in control and *LINC01179* knockdown 22Rv1 cells. (**C**) qRT-PCR analysis of the mRNA levels of *TLL1* in control and *LINC01179* overexpressed 22Rv1 cells. (**D**) Western blotting analysis of TLL1 expression levels in control and *LINC01179* overexpressed 22Rv1 cells. (**E**) Subcellular distribution was performed to isolate cytoplasmic and nucleoplasmic associated RNA. qRT-PCR was utilized to determine the subcellular localization ratio of *LINC01179* transcripts. *Actin* and *U6* were used as marker RNAs for quality control. (**F**) Representative images showing localization of *LINC01179* RNA transcripts (red) within nuclei of PC3 cells. Scale bar = 20 μm. (**G**) Silver staining of *LINC01179* RNA pull-down proteins. Mass spectrometry analysis of proteins that interacts with *LINC01179*. Anti-sense probe for *LINC01179* was used as control. (**H**) *LINC01179* RNA pull-down followed by western blot validated the interaction with Miz1. (**I**) The potential regulatory transcription factors for TLL1. (**J**) ChIP-qPCR results showed Miz1 chromatin binding at *TLL1* intron 1 region in control and *LINC01179* overexpressed PCa cells. (**K**) The luciferase reporter assay showed Miz1 reversing the increased effect of enhancer to the activity of *TLL1* promoter. (**L**) Pearson correlations between the expression of *TLL1* and *Miz1* in TCGA cohort. (**M**) Pearson correlations between the expression of *TLL1* and *Miz1* in DKFZ cohort. (**N**) qRT-PCR analysis of mRNA levels of *TLL1* and *Miz1* in control and Miz1 knockdown PC3 cells. (**O**) Western blotting analysis of TLL1 and Miz1 expression levels in control and Miz1 knockdown PC3 cells. (**P**) Western blotting analysis of TLL1 and Miz1 protein levels in control and Miz1 overexpressed PC3 cells. Data are shown as mean ± SD with three times of independent assay. NS, not significant, \*\**p* < 0.01, \*\*\**p* < 0.001. *p* values were assessed using two-tailed Student’s *t* tests.

### *LINC01179* attenuated PCa cells proliferation and migration by suppressing TLL1 expression and deactivating TGF-β signaling pathway

To date, the function of long-non coding RNA *LINC01179* has never been studied. We next investigated the functional role of *LINC01179* in prostate cancer. CCK8 assay showed that reduced expression of *LINC01179* apparently enhanced PCa cells growth (Fig. 4A, and fig. S4A). Colony formation assay revealed that knockdown of *LINC01179* in PCa cells promoted cells’ colony formation ability (Fig. 4, B and C, and fig. S4, B and C). Then, the effect of *LINC01179* on PCa cell migration ability was assessed by trans-well assay. The results indicated that *LINC01179* suppression in PCa cell lines significantly strengthened cells migration ability (Fig. 4, D and E, and fig. S4, D and E); increased expression of *LINC01179* weakened cells migration ability (Fig. 4, F and G, and fig. S4, F and G). Given that TLL1 could activate TGF-β signaling pathway to promote the PCa progression, we further explored the correlation between *LINC01179* and TGF-β signaling pathway. We found that contrary to TLL1, *LINC01179* inhibition in PC3 cells caused the increased phosphorylated smad2 levels (Fig. 4H); accordingly, *LINC01179* overexpression decreased phosphorylated smad2 levels (Fig. 4I). To further validate the relationship between *LINC01179* and TGF-β signaling pathway, we used TGF-β inhibitor (LY-2109761) to evaluate whether it could reverse the impact of *LINC01179* on PCa cells migration. The functional assays proved that LY-2109761 could significantly inhibit the increased migration ability of PCa cells with shRNA against *LINC01179* (Fig. 4, J to M). We further examined whether *LINC01179* suppressed PCa cells proliferation and migration by regulating TLL1 expression. We then knocked down TLL1 in *LINC01179*-knockdown PCa cells by infecting TLL1 knockdown lentivirus. As expected, re-knockdown of TLL1 rescued the increased PC3 cells proliferation ability caused by *LINC01179* suppression (Fig. 4N). Moreover, wound-healing assay suggested that *LINC01179* depletion increased PCa cells migration ability compared with control cells, and the enhanced migration ability was blocked after TLL1 knockdown (Fig. 4, O and P, and fig. S4, H and I). To further determine the tumorigenesis-suppressing role of *LINC01179* in vivo, subcutaneous xenograft mouse models were established. Stable *LINC01179*/TLL1 double knockdown PC3 cells, *LINC01179* knockdown PC3 cells or control cells were injected subcutaneously into male nude mice, respectively. One week after inoculation, tumors were measured every three days. As shown in Fig. 4, Q to S, *LINC01179* knockdown significantly inhibited tumor growth, reduced tumor volume and tumor weight compared to the control group, which was effectively reversed after TLL1 depletion. Altogether, our results suggested that *LINC01179* attenuated PCa cells proliferation and migration by suppressing TLL1 expression and deactivating TGF-β signaling pathway.

**Fig. 4.**
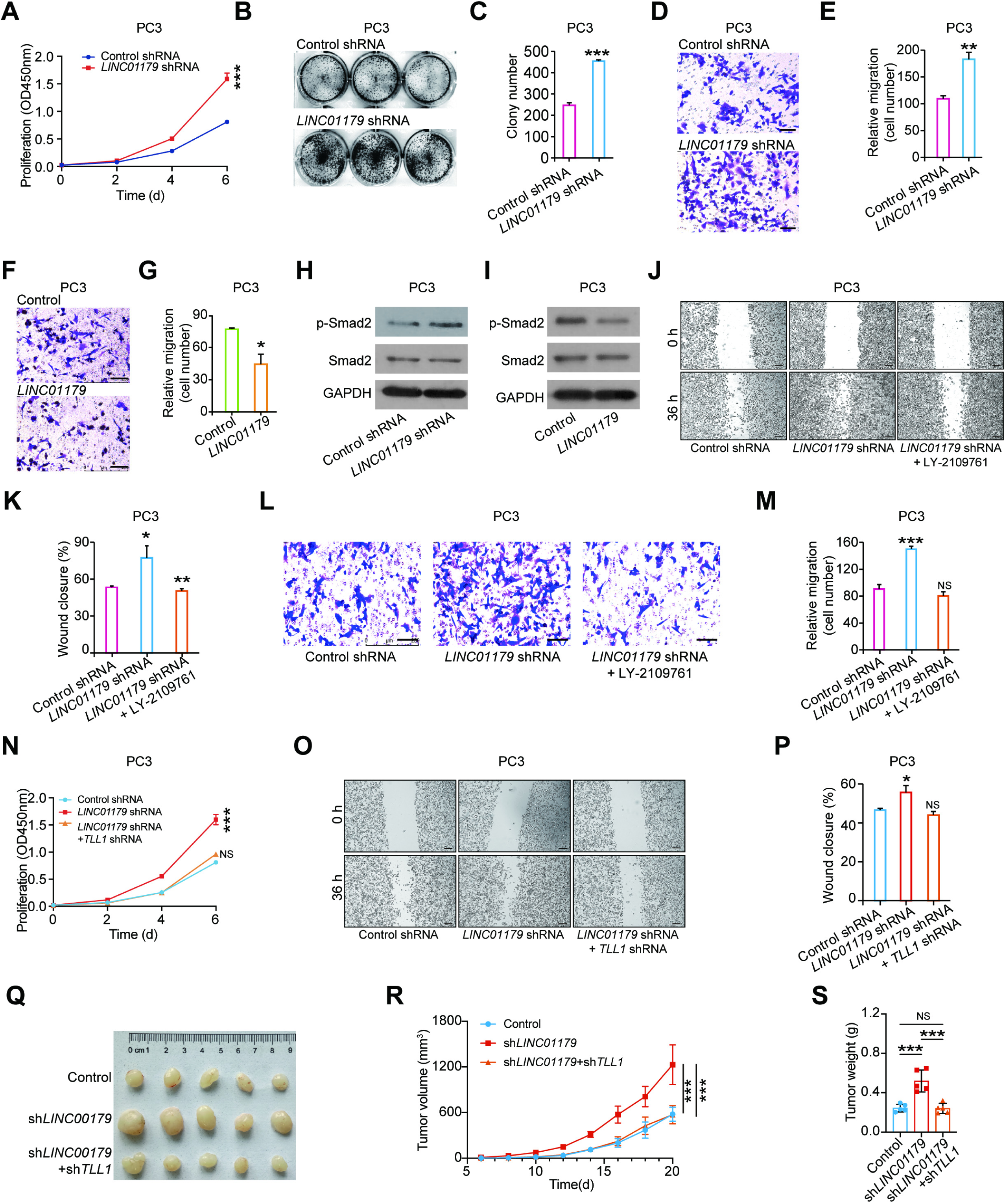
LINC01179 attenuated PCa cells proliferation and migration by decreasing TLL1 expression and deactivating TGF-β signaling pathway. (**A**) Effects of *LINC01179* suppression on cell proliferation measured by CCK8 assays in PC3 cells. (**B** and **C**) Representative pictures (B) and quantification analysis (C) of colony formation assays in control and *LINC01179* knockdown PC3 cells. (**D** and **E**) Representative pictures (D) and quantification analysis (E) of migration assays in control and *LINC01179* depleted PC3 cells. Scale bar = 100 μm. (**F** and **G**) Representative pictures (F) and quantification analysis (G) of migration assays in *LINC01179* overexpressed PC3 cells. Scale bar = 100 μm. (**H**) Western blotting analysis of p-Smad2 and Smad2 expression levels in control and *LINC01179* knockdown PC3 cells. (**I**) Western blotting analysis of p-Smad2 and Smad2 expression levels in control and *LINC01179* overexpressed PC3 cells. (**J** and **K**) Representative pictures (J) and quantification analysis (K) of wound healing assays in control, *LINC01179* knockdown and *LINC01179* knockdown treated with LY-2109761 (2 μM) PC3 cells. Scale bar = 200 μm. (**L** and **M**) Representative pictures (L) and quantification analysis (M) of migration assays in control, *LINC01179* knockdown and *LINC01179* knockdown with LY-2109761 (2 μM) treatment PC3 cells. Scale bar = 100 μm. (**N**) Effects of *LINC01179* knockdown and *LINC01179/*TLL1 double knockdown on cell proliferation measured by CCK8 assays in PC3 cells. (**O** and **P**) Representative pictures (O) and quantification analysis (P) of wound healing assays in control, *LINC01179* knockdown and *LINC01179/*TLL1 double knockdown PC3 cells. Scale bar = 200 μm. (**Q** to **S**) The control, *LINC01179* knockdown and *LINC01179/*TLL1 double knockdown PC3 cells were subcutaneously injected into BALB/c nude mice for xenograft growth (n = 5 per group). Tumor images (Q), tumor volumes (R) and tumor weights (S) are shown. NS, not significant, \**p* < 0.05, \*\**p* < 0.01, \*\*\**p* < 0.001. Data presented in **A**, **C**, **E**, **G**, **K**, **M**, **N** and **P** are shown as mean ± SD with three times of independent assay. In **A**, **C**, **E**, **G**, **K**, **M**, **N**, **P**, **R** and **S**, *p* values were assessed using two-tailed Student’s *t* tests.

### TLL1 regulated PD-L1 abundance in PCa cells through TGF-β signaling pathway

Considering that there is a close correlation between TGF-β signaling and PD-L1 [30], we proposed that TLL1 might regulate the expression of PD-L1. We first analyzed the expression correlation between *TLL1* and *PD-L1*, the results showed that *TLL1* is positively associated with *PD-L1* (Fig. 5, A and B, and fig. S5A). Then we inhibited TLL1 expression in PCa cells, RT-qPCR and western blot assays were performed to detect PD-L1 expression. The results indicated that TLL1 knockdown in PCa cells inhibited the mRNA and protein levels of PD-L1 (Fig. 5, C and D, and fig. S5, B and C). Moreover, suppressing TLL1 expression reduced cell surface PD-L1 expression in PCa cells detected by flow cytometry (Fig. 5, E and F, and fig. S5, D and E). Accordingly, ectopic expression of TLL1 led to the increased PD-L1 protein levels in PCa cells (Fig. 5G). To explore whether TLL1 regulates PD-L1 expression through TGF-β signaling pathway, TLL1 overexpressed PCa cells were treated with or without LY-2109761. The results showed that LY-2109761 could apparently decline the increased PD-L1 protein level caused by TLL1 overexpression (Fig. 5H and fig. S5F). In addition, we found that TLL1 knockdown can affect the mRNA levels of PD-L1 related ubiquitin enzymes (Fig. 5I and fig. S5G), including *OTUB1*, *USP5*, *USP22* and *SPOP* [31–36]. However, only *SPOP* mRNA level decreased after TLL1 was overexpressed in PCa cells (Fig. 5J and fig. S5H). Moreover, knockdown of TLL1 in PCa cells significantly enhanced SPOP protein levels and TLL1 overexpression resulted in the decreased SPOP protein levels (Fig. 5, K and L, and fig. S5, I and J). Besides, we found that the degradation of PD-L1 protein caused by TLL1 inhibition was obviously halted by the ubiquitination-proteasome pathway inhibitor MG132 (Fig. 5M and fig. S5K), suggesting that TLL1 enhanced PD-L1 protein stability through the ubiquitination-proteasome pathway. To further validate the results, we knocked down TLL1 in PCa cells and measured the endogenous ubiquitination of PD-L1 protein that was immunoprecipitated by the PD-L1 specific antibody. We found that TLL1 inhibition in PCa cells led to the increased ubiquitination of PD-L1 protein (Fig. 5N and fig. S5L). It was reported that the expression levels of SPOP were suppressed by TGF-β signaling [37], thus, we tested whether the TGF-β signaling pathway inhibitor could reverse the effect of TLL1 on SPOP protein levels. Then, PCa cells infected with TLL1 overexpression or control lentivirus were treated with or without LY-2109761 for 24 hours. Western blot assay indicated that the increased PD-L1 protein level caused by TLL1 overexpression in PCa cells was restrained by LY-2109761 treatment (Fig. 5O and fig. S5M). Collectively, these results proved that TLL1 suppressed SPOP expression and up-regulated the expression level of PD-L1 by activating TGF-β signaling pathway in PCa cells.

**Fig. 5.**
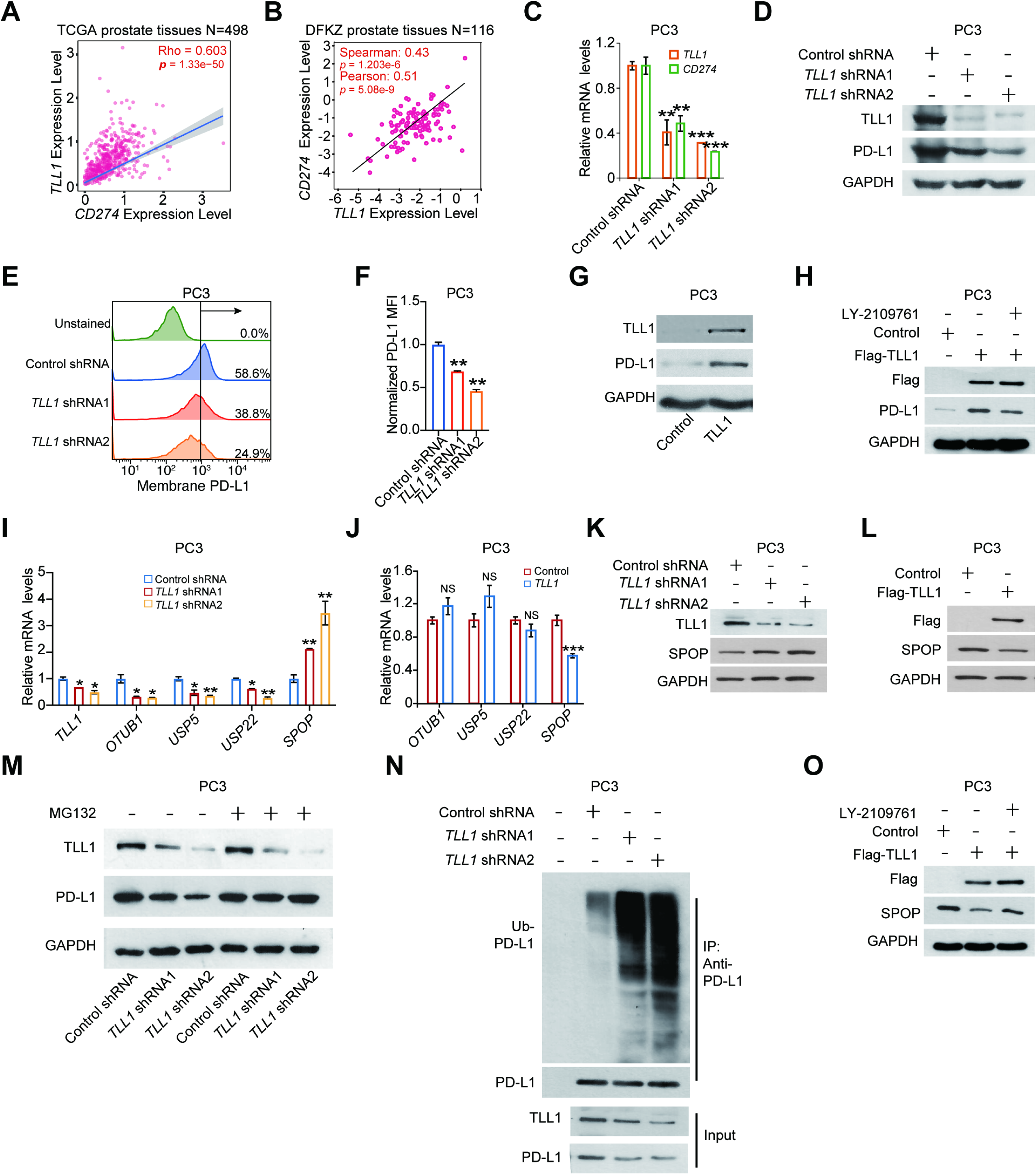
TLL1 suppressed SPOP expression and increased PD-L1 expression via activating TGF-β signaling pathway. (**A**) Pearson correlation between the expression of *TLL1* and *PD-L1* in TCGA cohort. (**B**) Pearson correlation between the expression levels of *TLL1* and *PD-L1* in DFKZ cohort. (**C**) qRT-PCR analysis of mRNA levels of *SPOP* in control and TLL1 knockdown PC3 cells. (**D**) Western blotting analysis of TLL1 and PD-L1 protein levels in control and TLL1 depleted PC3 cells. (**E**) Representative flow cytometry histograms analyzing cell surface PD-L1 in control and TLL1 knockdown PC3 cells are shown. (**F**) Bar graphs showed the fold change of cell surface PD-L1 in control and TLL1 knockdown PC3 cells as measured in (E). (**G**) Western blotting analysis of TLL1 and PD-L1 protein levels in control and TLL1 overexpressed PC3 cells. (**H**) Western blotting analysis of TLL1 and PD-L1 protein levels in control and TLL1 overexpressed PC3 cells treated with or without LY-2109761 (2 μM). (**I**) qRT-PCR analysis of mRNA levels of *TLL1*, *OTUB1*, *USP5*, *USP22* and *SPOP* in control and TLL1 knockdown PC3 cells. (**J**) qRT-PCR analysis of mRNA levels of *OTUB1*, *USP5*, *USP22* and *SPOP* in control and TLL1 overexpressed PC3 cells. (**K**) Western blotting analysis of SPOP protein levels in control and TLL1 knockdown PC3 cells. (**L**) Western blotting analysis of SPOP expression levels in control and TLL1 overexpressed PC3 cells. (**M**) Western blotting analysis of PD-L1 expression levels in TLL1 knockdown or control PC3 cells treated with or without MG132 (20 μM) for 8 hours. (**N**) The ubiquitination of PD-L1 was analyzed by immunoprecipitation in TLL1 knockdown PC3 cells and Western blot with indicated antibodies. Cell lysates were subjected to immunoblot with TLL1 or PD-L1 antibody. (**O**) PC3 cells transfected with Flag-TLL1 or control vector were treated with or without LY-2109761 (2 μM) for 24 hours. Cell lysates were subjected to immunoblot with Flag and SPOP antibody. NS, not significant, \**p* < 0.05, \*\**p* < 0.01, \*\*\**p* < 0.001. Data are presented as mean ± SD of triplicate experiments in **C**, **F**, **I** and **J**, *p* values were assessed using two-tailed Student’s *t* tests.

### TLL1 knockdown enhanced anti-tumor immunity in prostate cancer

Considering PD-L1 is an essential immune checkpoint protein, it binds to PD1 to evade T cell immunity, we explored whether TLL1 has any effects on T cell activity. Bioinformatics analysis in clinical PCa tumor samples showed that TLL1 was associated with CD8^+^ T cells and CD4^+^ T cells (Fig. 6A). Then, RM-1 cells were transfected with TLL1 knockdown lentivirus or control lentivirus, western blot assay was performed to detect TLL1 and PD-L1 expression levels. The results indicated that Tll1 suppression in RM-1 cells significantly weakened PD-L1 protein levels (Fig. 6B). Flow cytometry results further confirmed that Tll1 knockdown led to the decreased cell surface PD-L1 levels (Fig. 6, C and D). Subsequently, the Tll1 knockdown RM-1 cells or control cells were subcutaneously injected into mice, respectively, and anti-PD-1 antibody was applied in vivo to treat mouse tumors (Fig. 6E). Compared to the control group, the tumor size, tumor volume and tumor weight in the shTll1 group were decreased. However, the control group treated with anti-PD-1 antibody showed no significant change in the tumor size, tumor volume and tumor weight compared with the control group without anti-PD-1 antibody. Expectedly, Tll1 knockdown apparently enhanced the antitumor efficacy by anti-PD-1 antibody (Fig. 6, F to H). Next, we analyzed the characteristics of tumor-infiltrating lymphocytes (TILs) by flow cytometry and found that the infiltration proportions of CD3^+^ T cells and CD3^+^CD8^+^ T cells rather than CD3^+^CD4^+^ T cells were significantly increased in Tll1 knockdown group treated with anti-PD-1 antibody (Fig. 6, I to L). Moreover, IHC analysis of tumor tissues demonstrated that the PD-L1 abundance was dramatically reduced in Tll1 knockdown group, shTll1group does not differ significantly compared to shTll1 group treated with anti-PD-1 antibody (Fig. 6M). The proliferating index of tumor cells was reduced in shTll1 group or shTll1 group treated with anti-PD-1 antibody as determined by immunostaining of Ki67 (Fig. 6N). And p-Smad2 level of tumor cells was decreased in shTll1 group or shTll1 group treated with anti-PD-1 antibody (Fig. 6O). Taken together, these results indicated that TLL1 knockdown could enhance the antitumor efficacy by anti-PD-1 antibody via augmenting the infiltration proportions of CD8^+^ T cells.

**Fig. 6.**
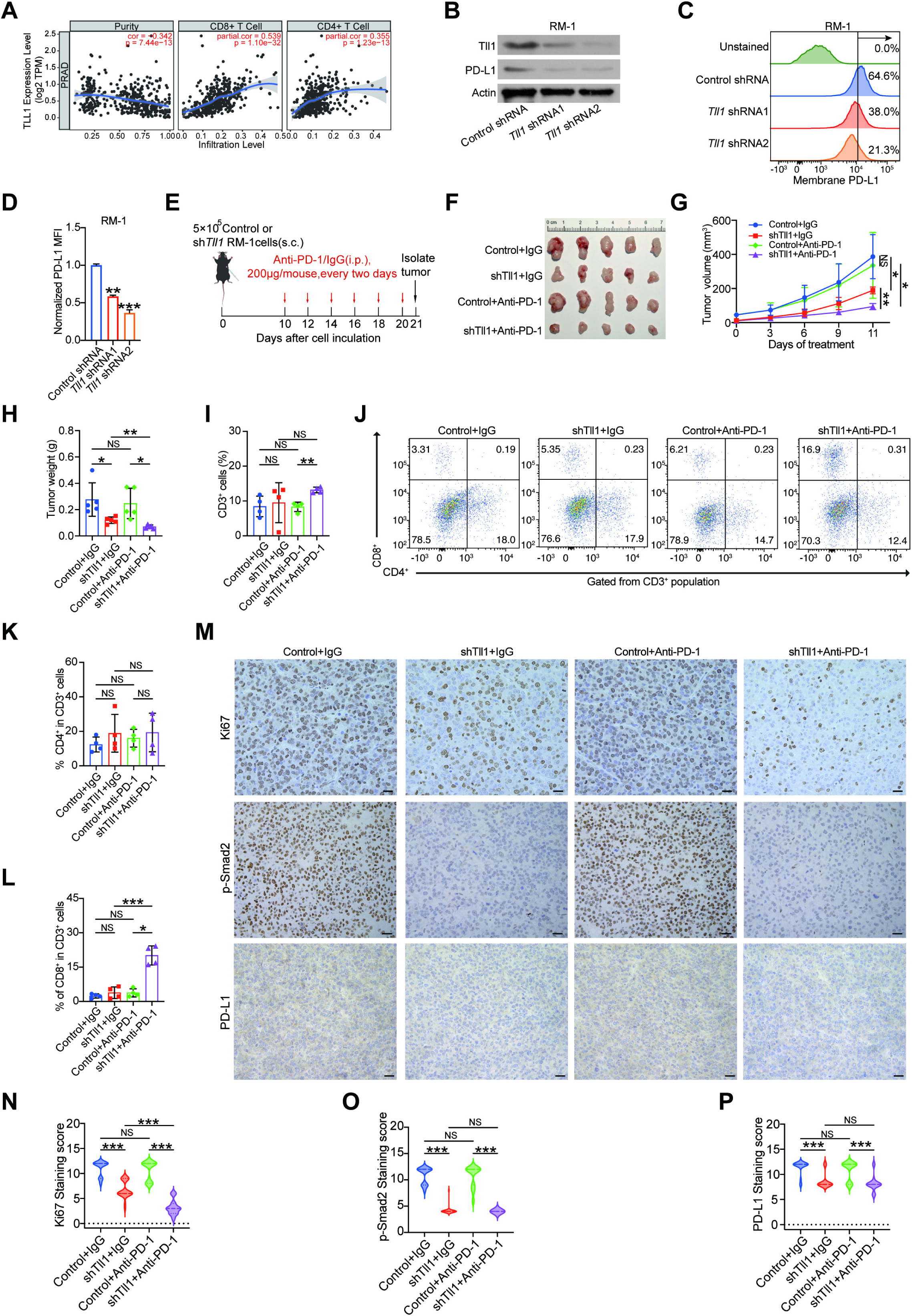
TLL1 knockdown enhanced anti-tumor immunity in prostate cancer. (**A**) The correlations between TLL1 expression and the immune cell infiltration (TIMER) in prostate cancer. (**B**) Western blotting analysis of TLL1 and PD-L1 protein levels in control and TLL1 depleted RM-1 cells. (**C**) Representative flow cytometry histograms analyzing cell surface PD-L1 in control and Tll1 knockdown RM-1 cells are shown. (**D**) Bar graphs showed the fold change of cell surface PD-L1 in control and Tll1 knockdown RM-1 cells as measured in (C). (**E**) The schematic of tumor inoculation and treatment in C57BL/6J mice. (**F** to **H**) Photographs (F), tumor growth curves (G) and tumor weight (H) of implanted tumors of RM-1 cells with Tll1 knockdown and control cells after inoculation in C57BL/6J mice. (**I**) Bar graphs of CD3^+^ T cells distribution in tumor tissue. (**J**) CD3^+^CD4^+^ and CD3^+^CD8^+^ T cells derived from tumor tissue were quantified by flow cytometry. (**K** and **L**) Quantitation of the percentages of CD3^+^CD4^+^ (K) and CD3^+^CD8^+^ (L) T cells in tumor tissue. (**M**) Representative images of H&E staining and IHC of Ki67, p-Smad2 and PD-L1of the subcutaneous tumors. Scale bar = 20 µm. (**N** to **P**) Ki67 (N), p-Smad2 (O) and PD-L1 (P) staining score in different groups. NS, not significant, \**p* < 0.05, \*\**p* < 0.01, \*\*\**p* < 0.001. Data are presented as mean ± SD of triplicate experiments in **D**. In **D**, **G**, **H**, **I**, **K**, **L**, **N**, **O** and **P**, *p* values were assessed using two-tailed Student’s *t* tests.

### TLL1 overexpression in T cells reduced the percentage of CD8^+^ T cells

To investigate the function of TLL1 in T cells, we generated *Tll1^tg^/^tg^*mice, in which a minigene consisting of a CAG promoter, a loxP-STOP-loxP (LSL) cassette, and *Tll1* cDNA is knocked into the Rosa26 gene locus. *Tll1^tg^/^tg^* mice were crossed to a transgenic mouse expressing Cre recombinase under the control of the mouse lymphocyte protein tyrosine kinase (Lck) promotor to generate *Tll1^tg/tg^*Lck-Cre^+^ mice (Fig. 7A). This promotor is active during the embryonic stages of thymic development and in mature T cells.

**Fig. 7.**
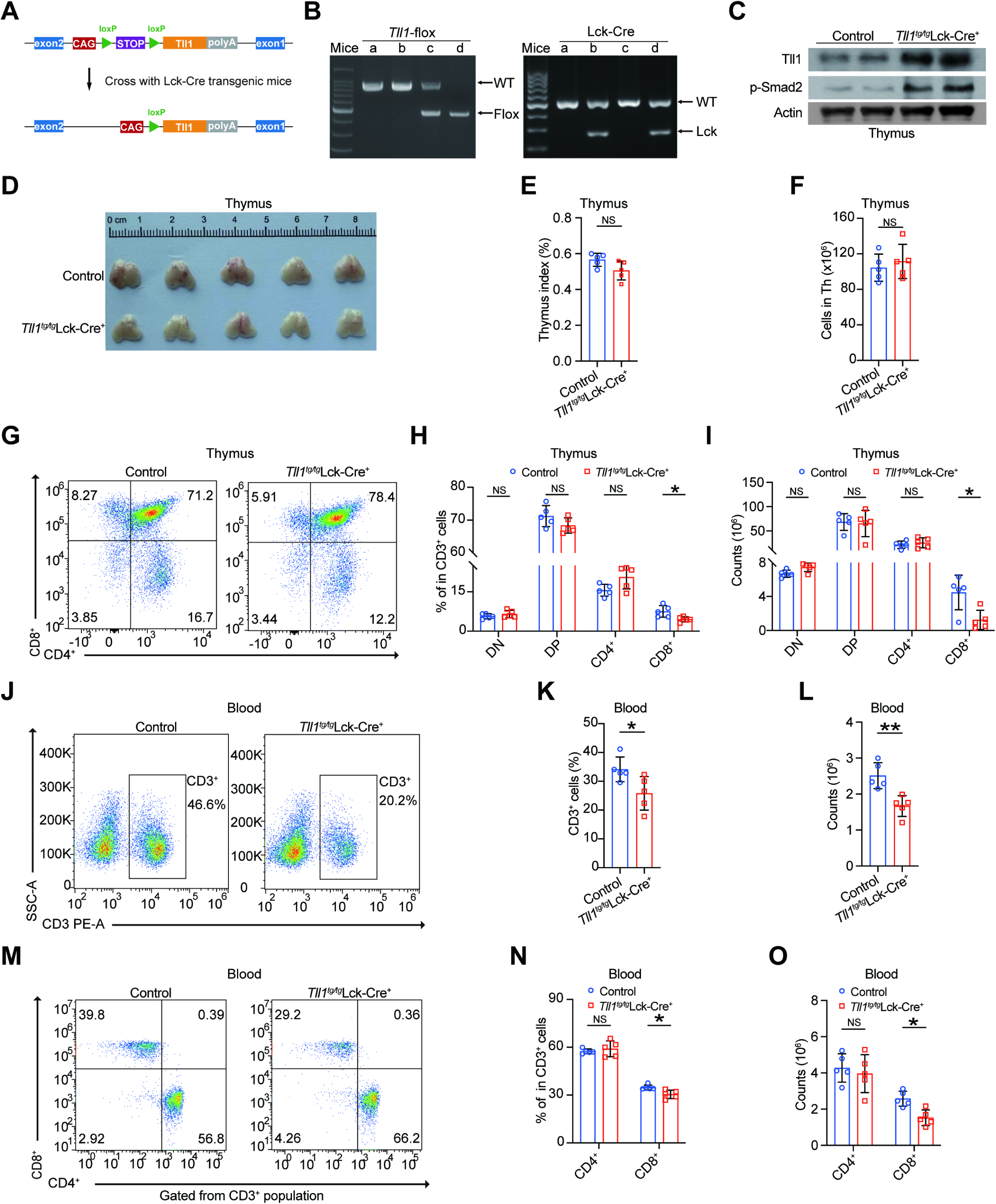
TLL1 overexpression in T cells reduced the percentage of CD8^+^ T cells. (**A**) Schematic diagram of T cell-specific *Tll1*-overexpressing mice. The mouse *Tll1* open reading frame was targeted to the ROSA26 locus preceded by a floxed (fl) transcriptional stop cassette and overexpressed by Cre-mediated recombination. (**B**) Genotyping: The Tll1 floxed allele (463 bp) was detected using primers P1 and P2, and the Lck-Cre transgene (272 bp) was detected using Lck-Cre primers. Mouse d was a *Tll1^tg/tg^*Lck-Cre^+^ mouse. (**C**) Western blotting analysis of Tll1 and p-Smad2 protein levels in control and *Tll1^tg/tg^*Lck-Cre^+^ mice thymus tissue. (**D**) Gross appearance of the thymus of control (*Tll1^tg/tg^*) and *Tll1^tg/tg^*Lck-Cre^+^ mice (n = 5 per group). (**E**) Thymus index of control and *Tll1^tg/tg^*Lck-Cre^+^ mice. (**F**) Bar graphs of the absolute cell numbers for total thymocytes of control and *Tll1^tg/tg^*Lck-Cre^+^ mice. (**G**) Representative flow cytometric analysis of surface expression of CD4^+^ and CD8^+^ T cells in the thymus of control and *Tll1^tg/tg^*Lck-Cre^+^ mice. (**H**) Bar graphs of the percentages of DN (CD4^-^CD8^-^), DP (CD4^+^CD8^+^), CD4^+^ and CD8^+^ T cells in the thymus of control and *Tll1^tg/tg^*Lck-Cre^+^ mice. (**I**) Bar graphs of the total cell number of DN, DP, CD4^+^ and CD8^+^ T cells in the thymus of control and *Tll1^tg/tg^*Lck-Cre^+^ mice. (**J**) Representative flow cytometric analysis of surface expression of CD3^+^ T cells in peripheral blood of control and *Tll1^tg/tg^*Lck-Cre^+^ mice. (**K** and **L**) Bar graphs of the percentages (K) and total cell number (L) of CD3^+^ T cells in peripheral blood of control and *Tll1^tg/tg^*Lck-Cre^+^ mice. (**M**) Representative flow cytometric analysis of surface expression of CD3^+^CD4^+^ and CD3^+^CD8^+^ T cells in peripheral blood of control and *Tll1^tg/tg^*Lck-Cre^+^ mice. (**N** and **O**) Bar graphs of the percentages (N) and total cell number (O) of CD3^+^CD4^+^ and CD3^+^CD8^+^ T cells in peripheral blood of control and *Tll1^tg/tg^*Lck-Cre^+^ mice. NS, not significant, \**p* < 0.05, \*\**p* < 0.01. In **E**, **F**, **H**, **I**, **K**, **L**, **N** and **O**, *p* values were assessed using two-tailed Student’s *t* tests.

Genotyping (Fig. 7B) and western blot (Fig. 7C) analyses demonstrated the overexpression of TLL1 was in thymocytes. Then, we evaluated the effect of TLL1 overexpression on thymocyte development. The weight and cell number of the thymus and spleen were normal in the *Tll1^tg/tg^*Lck-Cre^+^ mice (Fig. 7, D to F). The proportion and number of different T-cell subsets were also analyzed in the thymus of *Tll1^tg/tg^*Lck-Cre^+^ mice. The results showed that the proportion and number of double negative (DN) (CD4^-^CD8^-^), double positive (DP) (CD4^+^CD8^+^) and CD4^+^ cells were unaltered in *Tll1^tg/tg^*Lck-Cre^+^ mice. However, the proportion and number of CD8^+^ T cells was significantly decreased in *Tll1^tg/tg^*Lck-Cre^+^ mice compared to their control littermates (*Tll1^tg/tg^*) (Fig. 7, G to I).

To investigate whether the disruption in T cell development in thymus from *Tll1^tg/tg^*Lck-Cre^+^ mice has consequences for T cell populations in peripheral lymphoid organs, we analyzed the T cell populations in peripheral blood (PB), spleen and lymph nodes (LNs) from *Tll1^tg/tg^*Lck-Cre^+^ and littermate control mice. The results showed that the proportion and number of peripheral blood CD3^+^ T cells were decreased in *Tll1^tg/tg^*Lck-Cre^+^ mice compared to their control littermates (Fig. 7, J to L). Further, the proportion and number of peripheral blood CD3^+^CD4^+^ T cells were unaltered and the proportion and number of CD3^+^CD8^+^ T cells were reduced in *Tll1^tg/tg^*Lck-Cre^+^ mice (Fig. 7, M to O). The proportion and number of different T-cell subsets were also normal in the spleen of *Tll1^tg/tg^*Lck-Cre^+^ mice (fig. S6, A and D). The composition of T cells in lymph nodes was also unaltered *Tll1^tg/tg^*Lck-Cre^+^ mice (fig. S6, E to H). Collectively, these results showed that T cell specific overexpression of TLL1 disrupts T cell development in the thymus, which is associated with a decrease in the number and frequency of CD8^+^ T cells.

### TLL1 overexpression in T cells promoted RM-1 prostate tumor growth in mice by decreasing the numbers of tumor-infiltrating T cells

To further investigate the function of TLL1-overexpression T cells in vivo in the context of tumor immunity, we injected mouse RM-1 cells subcutaneously into *Tll1^tg/tg^*Lck-Cre^+^ and control mice, respectively. As shown in Fig. 8, A to C, tumors grew faster in *Tll1^tg/tg^*Lck-Cre^+^ mice and showed increased size and weight compared to those developing in control mice. On day 21 after injection of RM-1 cells, mice tumors and lymphocytes were isolated, counted and analyzed by FACS staining. Importantly, the percentage of CD3^+^ T cells in the tumors harvested from *Tll1^tg/tg^*Lck-Cre^+^ mice was significantly decreased compared to those from control mice (Fig. 8D). And the numbers of CD4^+^ T cells subsets were unaltered and CD8^+^ T cells subsets were decreased among the tumor infiltrating lymphocytes in *Tll1^tg/tg^*Lck-Cre^+^ mice (Fig. 8, E to G). Moreover, the expression of PD-1 in CD8^+^ T cells was increased in the *Tll1^tg/tg^*Lck-Cre^+^ mice compared with control mice (Fig. 8H). Moreover, we found that the weight and cells number of the thymus were normal in the *Tll1^tg/tg^*Lck-Cre^+^ mice (Fig. 8, I to K). The proportion and number of different T-cell subsets were also analyzed. The results showed that the proportion and number of DN, DP and CD4^+^ T cells were unaltered in *Tll1^tg/tg^*Lck-Cre^+^ mice. However, the proportion and number of CD8^+^ T cells was decreased in *Tll1^tg/tg^*Lck-Cre^+^ mice compared to their control littermates (Fig. 8, L to N). Collectively, these data indicated that TLL1 overexpression in T cells promoted RM-1 prostate tumor growth in mice by decreasing the infiltration of CD8^+^ T cells into tumors.

**Fig. 8.**
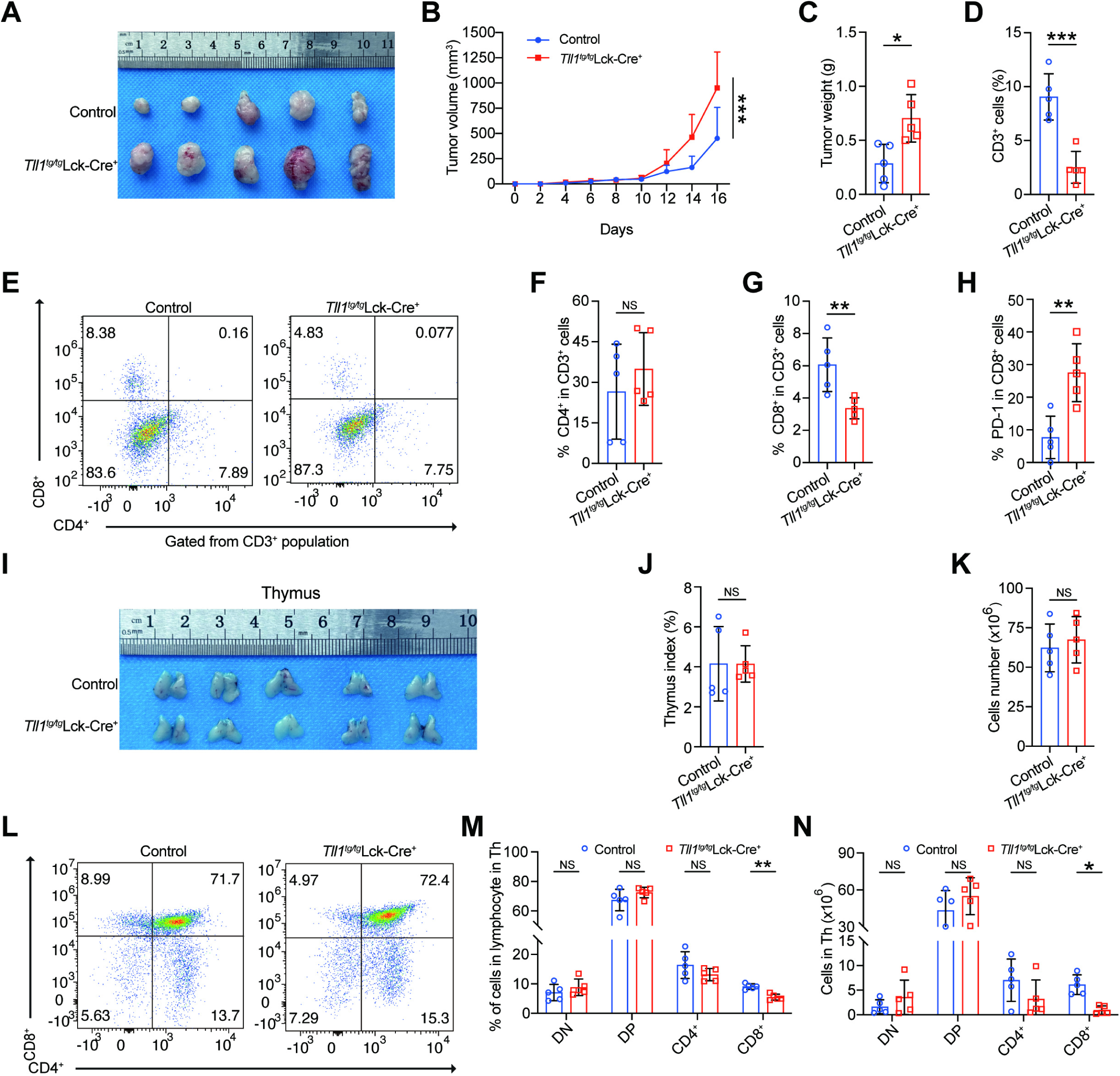
TLL1 overexpression in T cells promoted RM-1 prostate tumor growth in mice by decreasing the numbers of tumor-infiltrating T cells. (**A** to **C**) Tumor photographs (A), tumor growth curves (B) and tumor weight (C) of the control and *Tll1^tg/tg^*Lck-Cre^+^ mice after implanted tumors with RM-1 cells, n = 5 per group. (**D**) The percentages of CD3^+^ T cells in the tumors of control and *Tll1^tg/tg^*Lck-Cre^+^ mice. (**E**) Representative flow cytometric analysis of surface expression of CD3^+^CD4^+^ and CD3^+^CD8^+^ T cells in the tumors of control and *Tll1^tg/tg^*Lck-Cre^+^ mice. (**F** and **G**) Bar graphs of the percentage of CD3^+^CD4^+^ (F) and CD3^+^CD8^+^ (G) T cells in the tumors of control and *Tll1^tg/tg^*Lck-Cre^+^ mice. (**H**) Bar graphs of the percentage of PD-1^+^CD8^+^ T cells in the tumors of control and *Tll1^tg/tg^*Lck-Cre^+^ mice. (**I**) Gross appearance of the thymus of control and *Tll1^tg/tg^*Lck-Cre^+^ mice after implanted tumors with RM-1 cells. (**J**) Thymus index of control and *Tll1^tg/tg^*Lck-Cre^+^ mice after implanted tumors with RM-1 cells. (**K**) Bar graphs of the absolute cell numbers for total thymocytes of control and *Tll1^tg/tg^*Lck-Cre^+^ mice thymus after implanted tumors with RM-1 cells. (**L**) Representative flow cytometric analysis of surface expression of CD3^+^CD4^+^ and CD3^+^CD8^+^ T cells in the thymus of control and *Tll1^tg/tg^*Lck-Cre^+^ mice after implanted tumors with RM-1 cells. (**M**) Bar graphs of the percentages of DN, DP, CD4^+^ and CD8^+^ T cells in the thymus of control and *Tll1^tg/tg^*Lck-Cre^+^ mice after implanted tumors with RM-1 cells. (**N**) Bar graphs of the total cell number of DN, DP, CD4^+^ and CD8^+^ T cells in the thymus of control and *Tll1^tg/tg^*Lck-Cre^+^ mice after implanted tumors with RM-1 cells. NS, not significant, \**p* < 0.05, \*\**p* < 0.01, \*\*\**p* < 0.001. *p* values were assessed using two-tailed Student’s *t* tests.

**Fig. 9.**
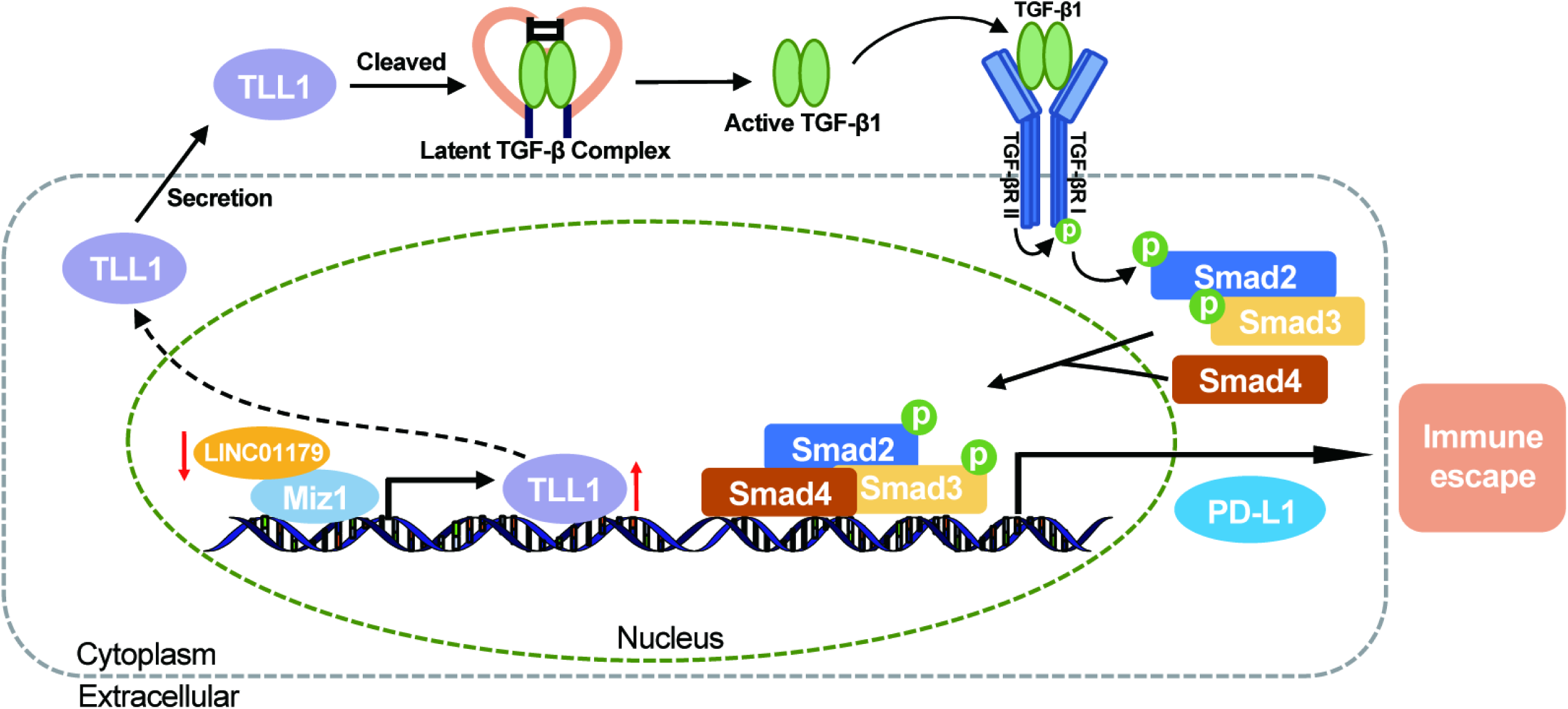
The mechanism underlying TLL1-mediated immune evasion of PCa cells. Proposed model for TLL1 promoted prostate cancer cells migration and metastasis through cleaving latent TGF-β1 to activate TGF-β signaling pathway. *LINC01179* interacted with Miz1 to attenuate TLL1 expression. Moreover, TLL1 increased the expression of PDL1 by activating TGF-β signaling pathway and TLL1 knockdown attenuates prostate cancer progression by enhancing antitumor immunity.

## DISCUSSION

In the present study, we found that metalloproteinase TLL1 was positively associated with prostate cancer aggressiveness. TLL1 promoted prostate cancer cells migration and metastasis via enhancing TGF-β signaling pathway. And TLL1 knockdown attenuates prostate cancer progression by enhancing antitumor immunity.

To date, the functional role of TLL1 in tumors has not been reported in the literature. Our study first showed that TLL1 was highly expressed in prostate tumors and positively associated with the poor prognosis of PCa patients. TLL1 depletion weakened cell growth and migration ability in PCa. Tumor xenograft models also indicated that TLL1 knockdown restrained tumor growth. Our findings suggested that TLL1 plays an oncogenic role in PCa progression.

It was reported that TLL1 could active TGF-β signaling pathway to control the differentiation of liver stem cells [27], however, how TLL1 activated the TGF-β signaling pathway is still unclear. In this study, we proved that TLL1 cleaved the latent TGF-β1 to activate TGF-β signaling pathway in prostate cancer cells. The activation of the TGF-β signaling pathway contributed to the progression of CRPC by triggering EMT [3]. Our findings showed that TLL1 suppression attenuated vimentin and enhanced E-cadherin protein levels. And the enhanced migration and metastasis of prostate cancer cells caused by TLL1 overexpression can be effectively inhibited by LY-2109761. Our study provides more clues to deeply understand the association between TLL1 and TGF-β signaling pathway in tumor cells.

LncRNAs have been found to play important role in tumor progression and are implicated in the modulation of gene expression [22]. We found that a novel lnc RNA *LINC01179* functions as an upstream regulator of TLL1 and modulates its expression in an inverse manner. It has been documented that lncRNAs regulate gene expression through multiple pathways at multiple levels [38,39]. Our study indicated that *LINC01179* can interact with transcription factor Miz1 to restrain the expression of TLL1. So far, the functional role of *LINC01179* in prostate cancer has not been reported. We found that *LINC01179* knockdown promoted prostate cancer cells growth and migration. It was reported that TGF-β signaling restrains the inhibitory function of MYC/Miz1 to the expression of p15 decades ago [24,40]. Our findings pointed out the suppressive effects of *LINC01179*/Miz1 on TGF-β signaling pathway in prostate cancer cells. The detailed regulatory mechanisms between *LINC01179* and Miz1 can be further explored to help us to better understand its function in prostate cancer.

The expression of immune checkpoint PD-L1 in tumor cells is crucial for their escape from antitumor immunity [41]. The increasing evidence demonstrated that the regulation of PD-L1 expression has been widely investigated at different levels, including transcriptional, translational, and post-translational levels, in various cancer types [42–45]. In the present study, we found that TLL1 positively regulates the mRNA and protein levels of PD-L1. TGF-β signaling pathway inhibitor could apparently diminish the increased PD-L1 protein level caused by TLL1. Moreover, TLL1 suppressed SPOP expression by activating TGF-β signaling pathway in PCa cells. SPOP, as a crucial E3 ubiquitin enzyme, can promote the ubiquitination of PD-L1 and degradation [46]. Our results also revealed that TLL1 inhibition accelerated the ubiquitination of PD-L1 and degradation, which was obviously halted by the ubiquitination-proteasome pathway inhibitor MG132. These results suggested that TLL1 might control the transcriptional level of PD-L1 and be also involved in regulating post-translational modifications of PD-L1 by TGF-β signaling pathway.

The binding of PD-1 to its ligand PD-L1 plays a crucial role in suppressing activated T-lymphocytes [47]. CD8^+^ T cells play critical roles in eliminating cancer cells [48]. And CD8^+^ T cells exhibit elevated expression of the inhibitory receptor PD-1, which is linked to the “exhausted” phenotype of CD8^+^ T cells [49]. In our study, we found that TLL1 knockdown could enhance the antitumor efficacy by anti-PD-1 antibody via augmenting the infiltration proportions of CD8^+^ T cells in vivo. Our study demonstrated that combining TLL1-shRNAs with anti-PD-1 antibody could enhance the antitumor response and offer a novel therapy to be tested in clinical trials with PCa patients.

Moreover, to investigate the function of TLL1 in T cells, we generated *Tll1^tg/tg^* mice. Upon crossing *Tll1^tg/tg^*mice with Lck-Cre^+^ mice, Cre recombinase-mediated activation excises the LSL cassette, allowing the CAG promoter to drive the constitutive expression of TLL1 in the T cells of thymus. We found that TLL1 overexpression in T cells impaired T cell development in the thymus. We then analyzed the proportion and number of different T-cell subsets in the thymus of *Tll1^tg/tg^*Lck-Cre^+^ mice. The results showed that the proportion and number of CD8^+^ T cells was significantly decreased in *Tll1^tg/tg^*Lck-Cre^+^ mice compared to their control littermates. And the proportion and number of CD3^+^CD8^+^ T cells was also reduced in *Tll1^tg/tg^*Lck-Cre^+^ mice.

In the tumor microenvironment, PD-L1, the ligand of PD-1, is expressed on the surface of tumor cells and binds to PD-1 on T cells, which, in turn, suppresses T cell effector functions [8,50]. In the present study, we observed that the percentage of CD3^+^ T cells in the tumors from *Tll1^tg/tg^*Lck-Cre^+^ mice was significantly decreased compared to those from control mice, probably because PD-L1 promotes T cell apoptosis when it binds to PD-1 expressed on the T cells [51]. And the numbers of CD4^+^ T cells subsets were unaltered and CD8^+^ T cells subsets were decreased among the tumor infiltrating lymphocytes in *Tll1^tg/tg^*Lck-Cre^+^ mice. Moreover, the expression of PD-1 in CD8^+^ T cells was increased in the *Tll1^tg/tg^*Lck-Cre^+^ mice compared with control mice, suggesting that TLL1-oversxpression CD8^+^ T cells are more exhausted within the tumor microenvironment. Tumors often exploit these negative feedback loops to establish a comprehensive immunosuppressive state, allowing them to evade the immunosurveillance against cancer [52,53]. Indeed, as presented in this study, the tumors that grew in *Tll1^tg/tg^*Lck-Cre^+^ mice have bigger size than in control mice. However, how TLL1 overexpression facilitates CD8^+^ T cells exhaustion need to be further investigated.

In summary, we found that TLL1 plays an oncogenic role in prostate cancer. And our study uncovers a novel role of TLL1 in modulating PD-L1 expression and reveals a novel therapeutic approach to improve the clinical efficacy of immunotherapy.

## MATERIALS AND METHODS

### Cell Lines and Cell Culture

Human embryonic kidney cells HEK293T, human prostate cancer cell lines (22Rv1, PC3), and mouse prostate cancer cell line (RM-1) were purchased from American Type Culture Collection (ATCC) and were tested negative for mycoplasma. 22Rv1 and RM-1 cells were cultured in RPMI 1640 Medium (Gibco, Cat#31800022). PC3 cells were cultured in Kaighn’s Modification of Ham’s F-12 Medium (HyClone, Cat#SH30526.01). HEK293T cells were cultured in DMEM (Gibco, Cat#12800017). All cultured media were supplemented with 10% fetal bovine serum (Gibco, Cat#10099-141C) and 1% penicillin/streptomycin (Solarbio, Cat#P1400). All cell lines were kept under controlled temperature (37°C) and CO2 (5%).

### Mice

Male BALB/c nude mice aged 4-5 weeks were acquired from Huafukang Biological Technology (Beijing, China) and maintained in an specific-pathogen-free (SPF) facility with a 12/12 h day/night cycle and unrestricted access to food and water. The Ethics Committee of the Laboratory Animal Management of Shaanxi Normal University gave its approval to this animal experiment (2023-038).

### Generation of T-cell-specific *Tll1*-overexpression mice

R26-LSL-*Tll1* and *Lck-Cre* transgenic C57BL/6J mice (Cat#CM-KI-231829) were purchased from Shanghai Model Organisms Center (Shanghai, China). To overexpress *Tll1* specifically in T cells, conditional *Tll1^tg/tg^* mice were crossed with mice expressing Cre recombinase under the control of the Lck proximal promoter to generate *Tll1^tg/tg^*Lck-Cre^+^ mice, and age-matched and sex-matched *Tll1^tg/tg^*C57BL/6J mice were used as a control. To detect the *Tll1* transgene in *Tll1* Tg mice, genomic DNA was extracted from tails of WT (wild type), control (*Tll1^tg/tg^*) and *Tll1* Tg mice (*Tll1^tg/tg^*Lck-Cre^+^), and then PCR analysis of the *Tll1* transgene was performed using two pairs of primers P1 (Forward: 5′-TCAGATTCTTTTATAGGGGACACA-3′; Reverse: 5′-TAAAGGCCACTCAATGCTCACTAA-3′) And P2 (Forward: 5′-GTCAGGGCCACCAGAAGAAA-3′; Reverse: 5′-AAAGTCCCGGAAAGGAGCTG-3′), which yielded a 463 bp band for the flox allele and a 967 bp band for wild-type allele. The presence of the Lck-Cre was detected with primer pair P3 (Forward: 5′-AGCGATGGATTTCCGTCTCTGG-3′; Reverse: 5′-AGCTTGCATGATCTCCGGTATTGAA-3′), which generated a PCR product of 272 bp.

### Plasmids, Lentivirus Production and Infection

For gene overexpression experiments, full-length human TLL1, and Miz1 were amplified by PCR and inserted into pLVX-mCMV-Zsgreen1-puro vector with Flag-tag. Full-length *LINC01179* was commercially synthesized and inserted into pLVX-mCMV-Zsgreen1-puro vector. For knockdown experiments, TLL1, Miz1 and *LINC01179* targeting shRNA designed using shRNA sequence designer software and the oligonucleotide were commercially synthesized and cloned to the pLKO.1-TRC vector, respectively. Viral production was performed in the HEK293T cell line, after transfection using Lipofectamine 2000 Transfection Reagent (Thermo Fisher Scientific, Cat#11668030) in antibiotic-free DMEM medium. For second-generation lentiviral production, 1.5 μg shRNA vector, 0.375 μg pMD2.G (envelope plasmid) and 1.125 μg psPAX2 (packaging plasmid) were diluted with 125 µL Opti-MEM (Invitrogen, Cat#11058-021) and combined with 125 µL Opti-MEM containing 6 µL Lipofectamine 2000. Lentiviral supernatant was harvested from day 2 to day 4, filtered through a 0.45 μM filter unit and stored at -80°C. For viral infection, viral supernatant containing 4 μg/mL polybrene (Solarbio, Cat#H8761) was added to PCa cells that were seeded 24h before infection at 60%-70% confluency. For the lentivirus-mediated overexpression or knockdown experiment, virus was removed and replaced by normal medium after 24 h, after which cells were selected with 1 μg/mL puromycin (Solarbio, Cat#P8230). When the normal PCa cells added with puromycin were completely dead, the target cells were cultured in normal growth medium for further experiments.

### Western Blot

The western blot assay was performed as previously described [54]. Briefly, cells were harvested and lysed with ice-cold cell lysis buffer (20 mM Tris-HCl, pH7.4; 0.5 M NaCl; 1% NP-40; 5 mM EDTA, pH 8.0; 1mM DTT; protease Inhibitor; phosphatase inhibitor). Then, cell lysate was cleared by spin down sample at 12,000 g for 15 min at 4°C. Protein concentrations were quantified with BCA Protein Assay Kit (Thermo Fisher Scientific, Cat#23227). Protein lysates were boiled at 95°C for 15 min after mixing with 5×SDS loading buffer. Proteins were resolved by SDS-PAGE and then transferred to PVDF membranes (Pall Life Sciences, Cat#BSP0161). Membranes were blocked in 5% nonfat milk for 1 h at room temperature and then incubated with diluted primary antibody overnight at 4°C. The following day, membranes were washed 3 times with 1×TBST and incubated with horseradish peroxidase conjugated secondary antibodies for 1h at room temperature. After the third wash with 1×TBST, membranes were incubated with ECL Western Blotting Substrate (Thermo Fisher Scientific, Cat#32209) or SuperSignal West Pico PLUS Chemiluminescent Substrate (Thermo Fisher Scientific, Cat#34580) and properly developed to X-ray film in a dark room.

### Reagents and Antibodies

TGF-β receptor inhibitor LY-2109761 was purchased from MCE (Cat#HY-12075). Proteasome inhibitor MG-132 was purchased from TargetMol (Cat#T2154). Recombinant Human TGF-β1 was purchased from PeproTech (Cat#100-21). TLL1 Polyclonal Antibody (Invitrogen, Cat#PA5-24246); Miz1 Rabbit mAb (CST, Cat#14300); SPOP Rabbit mAb (HUABIO, Cat#HA721319); PD-L1 Rabbit mAb (HUABIO, Cat#ET1701-41); FLAG Tag Monoclonal Antibody (Sigma, Cat#F3165); MYC Monoclonal Antibody (Invitrogen, Cat#MA1-980); Phospho-SMAD2 Rabbit mAb (CST, Cat#3104); E-Cadherin Mouse mAb (CST, Cat #14472); Normal Rabbit IgG (CST, Cat#2729); Mouse Control IgG (ABclonal, Cat#AC011); Vimentin Rabbit mAb (CST, Cat#5741); Anti-TGF β1 antibody (Abcam, Cat#215715); β-Actin Mouse mAb (ABclonal, Cat#AC004); Smad2 Rabbit mAb (ABclonal, Cat#A7699); Ubiquitin Rabbit mAb (ABclonal, Cat#A19686); GAPDH Monoclonal antibody (Proteintech, Cat#60004-1-Ig); Goat anti-Mouse IgG (H+L) HRP Secondary Antibody (Invitrogen, Cat#31430); Goat anti-Rabbit IgG (H+L) HRP Secondary Antibody (Invitrogen, Cat#31460); Anti-mouse IgG for IP (HRP)(Abcam,Cat#ab131368); Mouse Anti-rabbit IgG mAb (HRP Conjugate) (CST, Cat#5127).

### Measurement of secreted TGF-β1

TLL1 knockdown or overexpression PC3 cells were cultured in 10 mL serum-free F-12 medium, respectively. After 48h, the supernatant was collected and centrifuged at 2000 rpm at 4°C for 5min to remove cell debris. Then, Amicon^®^ Ultra columns (Millipore, Cat#UFC9030) was used to concentrate the supernatant according to the manufacturer’s instructions. The concentrated supernatant was analyzed for TGF-β1 by western blotting.

### Total RNA Isolation and Quantitative RT-PCR

Total RNA extraction from cells was performed using Trizol (Takara, Cat#9109) according to the manufacturer’s instructions. Complementary DNA was synthesized using ABScript II cDNA First-Strand Synthesis Kit (ABclonal, Cat#RK20400). Quantitative reverse transcriptase polymerase chain reaction (qRT-PCR) was performed using SYBR Green qPCR Master Mix (TargetMol, Cat#C0006). Each gene was run in triplicate and normalized to Actin or GAPDH. Sequences of all primers are listed in **Supplementary Table 1**.

### Subcellular Distribution

To determine the cellular localization of *LINC01179*, cytoplasmic and nuclear RNA were isolated using PAKIS Kit (Thermo Fisher Scientific, Cat#AM1921) according to the manufacturer’s instructions. Extracted RNA was reverse-transcribed immediately, and the expression of genes was measured by qRT-PCR analysis. The U6 RNA was used as nuclear control and Actin mRNA as cytoplasmic control. The primers for target genes and the negative control were listed in **Supplementary Table 1**.

### RNA Fluorescence In Situ Hybridization (RNA-FISH)

The subcellular localization of *LINC01179* was analyzed using RNA fluorescence in situ hybridization. Briefly, 1.5×10^4^ PC3 cells were seeded on round glass coverslips in 12-well plate. After 24 h, cells were washed with PBS and fixed in 4% paraformaldehyde for 15 min at room temperature. Subsequently, cells were permeabilized in PBS containing 0.2% Triton X-100 for 40 min at room temperature and incubated in 2×SSC buffer for 30 min at room temperature after three washes with PBS for 5 min each. After that, the cells were washed with 70%, 85% and 100% ethanol for 3 min each time, respectively. Then, cells were incubated with 3µg 5’Cy3-labeled *LINC01179* probe in RNA-FISH Hybridization buffer (2×SSC; 10% Dextran sulfate; 0.2 mg/mL BSA; 10% Formamide; 40U/mL RNase inhibitor) at 37 °C overnight. On the following day, the cells were rinsed with 0.4×SSC/0.3% Tween and 2×SSC/0.1% Tween, respectively. After completing these steps, the nuclei were stained with DAPI (Roche, Cat#10236276001). The cells were visualized and photographed using Axio Imager Upright Microscope. The sequence of probe (5’ to 3’): Cy3-CAGTTCTCTGGTGAGATGCCAGGTAGCAGCGA.

### RNA-Pulldown Assay

The full-length *LINC01179* sense and antisense were PCR amplified using T7 promoter containing primer and then reversely transcribed and biotin-labeled by High Yield T7 Biotin16 RNA Labeling Kit (APExBio, Cat#K1082). Biotinylated RNA was folded in RNA structure buffer (10 mM Tris-HCl, pH 7.0; 0.1 M KCl; 10 mM MgCl2) at 90°C for 2 min, put on ice for 2 min immediately, and then transferred to room temperature for 30 min to allow proper RNA secondary structure formation. Biotinylated RNAs were mixed with 50 µL streptavidin-conjugated magnetic beads (Thermo Fisher Scientific, Cat#65001) in RIP buffer (150 mM KCl; 25 mM Tris-HCl, pH 7.4; 0.5 mM DTT; 0.5% NP-40; 1 mM PMSF; protease inhibitor) for 1 h at room temperature. Total cell lysate was freshly prepared and 1 mg of the cleared lysate was added to each binding reaction with protease inhibitor and RNase inhibitor. Then the mixture was incubated in Protein-RNA binding buffer (0.2 M Tris-HCl, pH 7.5; 0.5 M NaCl; 20 mM MgCl2; 1% Tween-20) with rotation for 2 h at 4 °C. After washing three times thoroughly, the RNA-protein binding mixture was boiled in 2×SDS loading buffer and the eluted proteins were resolved by SDS-PAGE followed by silver staining or mass spectrometry.

### Ubiquitination of PD-L1

The ubiquitination assay was performed as previously described [55]. Briefly, for detecting the endogenous ubiquitination of PD-L1, TLL1 knockdown PCa cells or control cells were lysed in lysis buffer, respectively. Then, the ubiquitination of PD-L1 was analyzed by immunoprecipitated with anti-PD-L1 antibody (HUABIO, Cat#ET1701-41) and immunoblotted with anti-Ub (ABclonal, Cat#A19686) antibody or anti-PD-L1 antibody.

### Chromatin Immunoprecipitation (ChIP)

The ChIP assay was performed as previously described [56]. Briefly, cells were cross-linked with a final concentration of 1% formaldehyde in growth medium for 10 min at room temperature, and cross-linking was quenched by the addition of glycine to a final concentration of 125 mM and incubation for 5 min at room temperature. Cells were collected from the dishes by scraping and washed three times with PBS. Cell pellets were resuspended in SDS lysis buffer (0.5% SDS; 10 mM EDTA; 50 mM Tris-HCl, pH 8.0) and sonicated to shear the chromatin to yield DNA fragment size at the average of 400 bp. Then, approximately 6μg of indicated antibodies or control IgG antibodies were incubated with 70 µL Dynabeads Protein G beads (Thermo Fisher Scientific, Cat#10004D) in IP buffer (2mM EDTA; 150mM NaCl; 20mM Tris-HCl, pH8.0; 1% Triton X-100; protease inhibitor) for 10 h before immunoprecipitation with the sonicated chromatin overnight at 4°C. After incubation, the complex was washed one time with Wash buffer I (20 mM Tris-HCl, pH 8.0; 2 mM EDTA; 0.1% SDS; 1% Triton X-100; 150 mM NaCl), Wash buffer II (20 mM Tris-HCl, pH 8.0; 2 mM EDTA; 0.1% SDS; 1% Triton X-100; 500 mM NaCl), Wash buffer III (10 mM Tris-HCl, pH 8.0; 1 mM EDTA; 250 mM LiCl; 1% Deoxycholate; 1% NP- 40). After rinsing with Wash buffer IV (10 mM Tris-HCl, pH 8.0; 1 mM EDTA) twice, precipitates were eluted in Extraction buffer (10 mM Tris-HCl, pH 8.0; 1 mM EDTA; 1% SDS) with 0.3 M NaCl and 1mg/mL proteinase K for 10-12 h at 65°C with gentle rocking. The reverse-crosslinked ChIP DNA samples were purified by MinElute PCR Purification Kit (Qiagen, Cat#28006). Quantification was performed using qRT-PCR with SYBR Green qPCR Master Mix. Control IgG and input DNA signal values were used to normalize the values. The primers for target genes and the negative control are listed in **Supplementary Table 1**.

### Cell Viability and Proliferation Assays

Cell viability and proliferation were detected by Cell Counting Kit-8 (CCK-8) (TargetMol, Cat#C0005) following the manufacturer’s instructions. In brief, control or treated cells were seeded into 96-well plates at a density of 2 × 10^3^ or 2.5 × 10^3^ cells per well. Subsequently, 10 μL CCK-8 reagent was added at specified time points. After incubating at 37°C for 4 h, optical density at 450 nm (OD450) was measured for each sample by a microplate reader (BioTek, USA). All experiments were conducted in triplicate. Significance was calculated by two-tailed Student’s t test.

### Migration Assay

For the migration assay, 3×10^4^ 22Rv1 cells or 2×10^4^ PC3 cells were resuspended completely with serum-free medium and added to the upper chambers of the Transwell plates (Corning, Cat#3422). At the same time, complete medium supplemented with 10% FBS was added to the lower chamber of the Transwell plates. After the indicated culture time, cells located on the bottom surface of the upper chambers were fixed with 3.7% formaldehyde, permeabilized with methanol, and stained with crystal violet. Migrant cells were counted in five random visual fields using an inverted Zeiss microscope. A two-tailed Student’s t test was used to perform statistical analysis from three replicate experiments.

### Wound Healing Assays

Wound healing assays were performed as follows. 2×10^5^ cells were seeded in a 12-well plate and scratched with a 200-µL pipette tip when 90-95% confluency was reached. Original scratch widths were recorded, and the scratch widths of cells after 24 h or 36 h were measured. The area of the wound in each well was analyzed using ImageJ software. The percentage of wound-healing was calculated using the following formula: (original scratch width - scratch width after healing) / (original scratch width) × 100%. All results are representative of three independent experiments. Statistical significance was calculated by the two-tailed Student’s t test.

### Colony Formation Assay

For the colony formation assay, cells were seeded at a density of 500 cells per well in 12-well plates and cultured for two weeks, with medium changed every 2-3 days. The colonies were treated with 3.7 % formaldehyde for 15 minutes. Following this, a stain of 0.1% crystal violet was applied for 30 minutes, and the colonies were counted using Image J software. All data came from at least three replicates, and statistical analysis was performed by two-tailed Student’s t test.

### Luciferase Reporter Assay

The TLL1 promoter was inserted into pGL3 Basic vector. Then, DNA fragments (TLL1 intron1) binding by Miz1 were added into the upstream of TLL1 promoter-pGL3 Basic vector. The internal Renilla control plasmid pGL4.75 [hRluc/CMV] (Promega), the target plasmids were reverse co-transfected into 22Rv1 cells using Lipofectamine 2000 Transfection Reagent according to the protocol provided by the manufacturer. The experiments were carried out on the 96-well white plates. 100 µL 2×10^5^ 22Rv1 cells/mL was added per well. After 48 h, luciferase activity was measured with Dual-Glo Luciferase Assay System (Promega, Cat#E2920). All data were obtained from at least three replicate wells and statistical analyses were performed with a two-tailed Student’s t test.

### Enzyme-Linked Immunosorbent Assay (ELISA)

The concentration of TGF-β1 in cell supernatant was detected by ELISA. When TLL1 overexpression or knockdown PCa cells grown to suitable density, the cell supernatant was centrifuged at 3000 rpm/min at 4°C for 10 min, and the content of TGF-β1 in the supernatant was detected by ELISA kit (JonlnBio, Cat#JL10706) according to the manufacturer’s instructions.

### Immunohistochemistry (IHC) and quantification

The PCa tissue chips (PRC1601) were purchased from Superbiotech (Shanghai, China) to explore the expression of TLL1 in PCa tissues and adjacent tissues, respectively. IHC staining was performed according to a standard streptavidin-biotin-peroxidase complex method. We first deparaffinized the tissue chips, and rehydrated using graded ethanol, rinsed with deionized water, and then blocked with 3% hydrogen peroxide for 30 min at room temperature. Antigen retrieval was performed by high-pressure-cooking the samples in a 10 mM citrate buffer (pH 6.0) for 5 min. Tissue chips were blocked with 5% BSA for 1 h at room temperature and subsequently incubated with TLL1 primary antibody (Invitrogen, Cat#PA5-24246) (diluted at 1:50) at 4 °C overnight. Then incubated the tissue chips with Envision-HRP secondary antibody for 1 h, the signals were detected by diaminobenzidine followed by hematoxylin counterstaining. Aperio ScanScope slide scanner was used for viewing the signals. Tumor tissue slides were deparaffinized, rehydrated, and treated with 3% hydrogen peroxide for 20 min at room temperature to block endogenous peroxidase activity. Slides were then subjected to antigen retrieval in EDTA buffer (pH 9.0) at 96 °C for 20 min. After blocking with 10% goat serum, slides were incubated with primary antibodies at 4 °C overnight. The following antibodies were used: Phospho-Smad2^S465/467^ Rabbit Polyclonal antibody (Abclonal, Cat#AP0548), PD-L1/CD274 Polyclonal antibody (Proteintech,Cat#17952-1-AP), Anti-Ki67 Rabbit pAb (Servicebio, Cat#GB111499). Following incubation with secondary antibodies and development with 3,3-diaminobenzidine (DAB), the final IHC score was calculated based on staining extent and intensity. For evaluation of immunohistochemical results, the intensity of staining was scored as follows: 0 (no staining), 1 (weak), 2 (moderate), and 3 (high). The percentage of immunolabeling was categorized into five groups: 0 (0%), 1 (1%–25%), 2 (26%–50%), 3 (51%–75%), 4 (76%–100%) and evaluated by two independent experienced pathologists. The product of the staining intensity multiplied by the percentage of immunolabeling was determined for each tumor sample. Finally, the final staining score was calculated and analyzed.

### H&E Staining

Murine lung and tumor tissue were fixed in 4% paraformaldehyde at room temperature. After 12 h, the tissues were dehydrated in an ascending gradient of ethanol and then made transparent with xylene. Subsequently, the lungs have been embedded in paraffin blocks and cut into sections of 3 μm thickness onto microscopic slides. For visualization of metastases, the sections were then stained for hematoxylin and eosin. For that, a rehydration step with a series of xylol (I), xylol (II) (each for 2 min), 100%, 96%, and 70% ethanol (for 1 min) has been performed. Samples were briefly rinsed with cold water (30 s), stained with hematoxylin for 6 min, rinsed with warm water for 4 min and with 70% ethanol for 1 min, stained with eosin for 2.5 min and dehydrated with a series of 96% (I), 96% (II), 96% (III), and 100% ethanol (each 1 min). Lastly, the slides were put into xylol (2.5 min). Finally, the sections were sealed with neutral gelatin after drying, and images were captured by the microscope system.

### Detection of cancer cell surface PD-L1 via flow cytometry

The expression of PD-L1 on cancer cell surface was detected by flow cytometric analysis. Briefly, 22Rv1, PC3, and RM-1 cells with stable knockdown of TLL1 and control cells were collected and washed twice with PBS. Then the cells were suspended in 2% FBS-PBS and stained with CD274 (PD-L1) Monoclonal Antibody (Thermo Fisher Scientific, Cat#25-5982-80) on 4°C for 30 min. After washing with PBS, the cells were resuspended in 2% FBS-PBS and detected by BD FACS Aria III (BD Biosciences, USA) flow cytometer. Data were analyzed by FlowJo software.

### Animal experiments

#### *in vivo* xenograft tumor models

(1) 1 × 10^6^ control, shTLL1, sh*LINC01179*, and sh*LINC01179* + shTLL1 PC3 cells were injected subcutaneously in 100 μL medium into the right flank of 4-5 weeks old BALB/c nude male mice (n = 5 per group). Tumor volume was measured with a caliper and after 20 days, mice were sacrificed and tumors were isolated and weighed. Tumor tissues were embedded in paraffin and dissected for H&E staining analysis. Tumor volume = length × width^2^ × 1/2.

(2) 5 × 10^5^ control or shTll1 RM-1 cells were injected subcutaneously in 100 μL medium into the right flank of mice. When the tumor volume reached roughly 50∼100 mm^3^, mice were pooled and randomly divided into 4 groups (control+IgG, shTll1+IgG, control+anti-PD-1, and shTll1+anti-PD-1, n=5 per group) with comparable average tumor size. Anti-PD-1 (MedChemExpress,Cat#HY-P99144; 200 μg/mouse) or control IgG (MedChemExpress,Cat# HY-P990679; 200 μg/mouse) was injected intraperitoneally into each mouse every two days. Tumor volumes were measured every 3 days using a digital caliper. When tumor volume reached about 500∼600 mm^3^, mice were sacrificed and tumors were isolated and weighed. The cells from the tumors were isolated after lysing red blood cells for subsequent flow cytometry analysis. Tumor tissues were embedded in paraffin and dissected for H&E and IHC analysis.

(3) 1 × 10^6^ RM-1 cells were injected subcutaneously in 100 μL medium into the right flank of 4-5 weeks old control and *Tll1^tg/tg^*Lck-Cre^+^ mice (n = 5 per group). Tumor volume was measured with a caliper every two days. When tumor volume reached about 1500 mm^3^, mice were sacrificed and tumors were isolated and weighed. The cells from the tumor and thymus were isolated after lysing red blood cells for subsequent flow cytometry analysis.

### *in vivo* lung metastasis models

15 male BALB/c nude mice at 4-5 weeks of age were randomly divided into three groups (n=5 per group). Then 1×10^6^ PC3-control cells (n=5) and PC3-TLL1 cells (n=10) were injected into the tail vein, respectively. 7 days later, 5 mice injected with PC3-TLL1 cells were orally treated with LY-2109761 (50 mg/kg every two days). 30 days later, lungs were dissected from each of the mice, and then H&E staining was used to examine the metastases status.

### Tumor Tissue Dissociation

Tumor tissues were collected from euthanized mice and washed twice with ice-cold PBS. The tissues were then cut into small pieces using sharp dissection scissors and incubated in 10 mL dissociation solution (0.2 mg/mL Collagenase D (Roche, Cat#11088866001); 0.05 mg/mL DNase I (Roche, Cat#10104159001)) for 1.5 h with continuous rotation at 37 °C. The digestion was quenched by adding 10 mL RPMI 1640 medium with 10% FBS and the cell suspension was passed through 70 µm cell strainer. After washing once with 2% FBS-PBS, cells were further separated using percoll (Cytiva, Cat#17-5445-01) gradient (70%/30%, v/v) centrifugation. After centrifugation at 500 g at 4°C for 25 min, the interphase was collected, and the single-cell suspension was washed once with PBS. After that, red blood cell lysis solution (Roche, Cat#11814389001) was added to resuspend the cells, and incubated at room temperature for 3 min for full lysis of red blood cells, followed by adding PBS to stop the lysis. After centrifugation of the lysis solution, the supernatant was discarded, and the cells were suspended in 2% FBS-PBS for further flow cytometry experiments.

### Flow Cytometry

Single cells isolated from mouse tumor, thymus, spleen, lymph node and blood were suspended in 100 µL 2% FBS-PBS, the antibodies were added according to manufacturer’s instructions, and incubated at 4°C for 30 min in dark. The following antibodies were used for flow cytometry: CD3e-PE (eBioscience, Cat#12-0031-81), CD4-FITC (eBioscience, Cat#11-0041-82), CD8a-APC (eBioscience, Cat#17-0081-81), CD279-PE (PD-1, eBioscience, Cat#12-9985-81). After washing with PBS, the cells were resuspended in 300 µL 2% FBS-PBS. All flow cytometry assays were performed on a BD FACS Aria III (BD Biosciences, USA) instrument. All data were analyzed using FlowJo software and Prism software. Statistical analyses were performed as indicated and *p* values of < 0.05 considered significant.

### Statistical Analysis

All of the statistical details including the statistical tests, exact number of animals, definition of center, and dispersion and precision measures for the experiments can be found in the figures, figure legends or in the results. All statistical analyses were performed by using R and GraphPad Prism 9.0. All data were presented as mean ± SD. Comparisons between two groups were performed by using an unpaired two-sided student’s *t*-test. For intravenous injection experiments and survival analysis, each mouse was counted as a biologically independent sample. For correlation analysis, we used the Spearman and Pearson coefficients. For survival studies, the Kaplan-Meier method was used to estimate and plot the survival curve, and the log rank test and Cox proportional hazard ratio analysis were used to evaluate differences in survival data among different groups. For all statistical tests, *p* < 0.05 was considered statistically significant. *Signifies *p* < 0.05, \*\**p* < 0.01, \*\*\**p* < 0.001. All experiments were repeated at least three times.

## AUTHOR CONTRIBUTIONS

P.G., X.M.D. and K.Z. conceived the project, J.L.H., J.Q.H., H.H., X.Y.W., Z.H.Z., X.Z., L.L.,Y.T.R., X.L. J.Y. and M.F. performed the experiments. J.L.H. and X.M.D. performed data analysis. P.G., X.M.D., K.Z. and J.L.H wrote the manuscript.

## COMPETING INTERESTS

The authors declare no competing interests.

## ETHICS APPROVAL

Animal experiments were performed according to the protocols approved by the ethical committee of Shaanxi Normal University (Approval number: 2023-038).

## ACKNOWLEDGEMENTS

This work was funded by the National Natural Science Foundation of China (82472665), Fundamental Research Funds for the Central Universities (GK202201004), Natural Science Foundation of Shaanxi Province (2023-JC-YB-716), Excellent Graduate Training Program of Shaanxi Normal University (LHRCTS23091) and College Students’ Innovative Entrepreneurial Training Plan Program (202410718037).

## ADDITIONAL INFORMATION

### Supplementary data

Supplementary Table 1. Sequences of primers and LINC01179 IP-MS data related to this article.

---

**Supplementary Figure 1.**
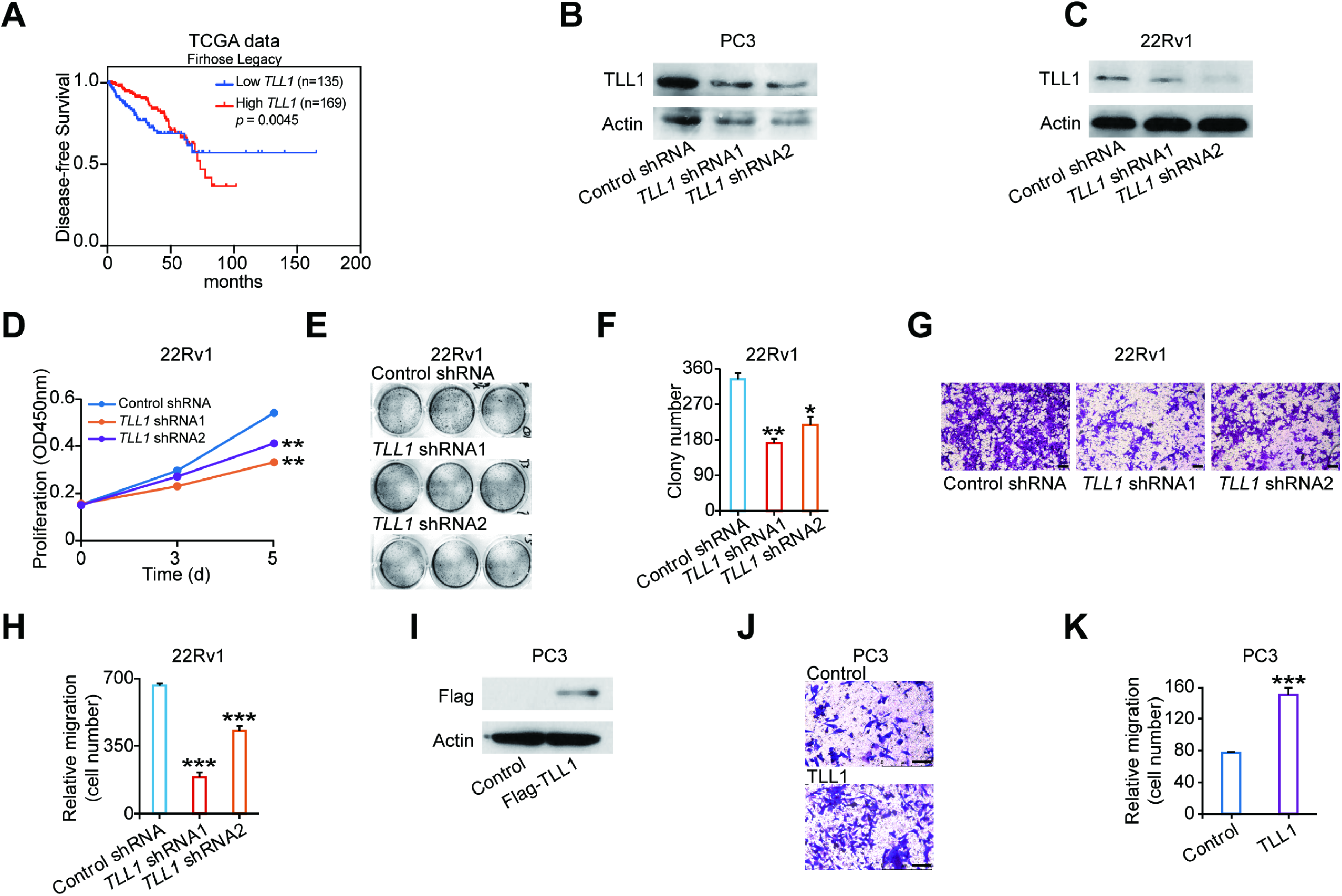
TLL1 knockdown weakened PCa cells growth and migration, related to Figure 1. (**A**) Association between Disease-free survival of prostate cancer patients and *TLL1* mRNA expression from the TCGA database. *p* values were assessed using log rank test. (**B**) Western blotting analysis of TLL1 protein levels in control and TLL1 depleted PC3 cells. GAPDH was used as a loading control. (**C**) Western blotting analysis of TLL1 protein levels in control and TLL1 depleted 22Rv1 cells. GAPDH was used as a loading control. (**D**) Effects of TLL1 knockdown on cell proliferation measured by CCK8 assays in 22Rv1 cells. (**E** and **F**) Representative pictures (E) and quantification analysis (F) of colony formation assays in control and TLL1 knockdown 22Rv1 cells. (**G** and **H**) Representative pictures (G) and quantification analysis (H) of migration assays in control or TLL1 knockdown 22Rv1 cells. Scale bar = 100 μm. (**I**) Western blotting analysis of TLL1 expression in control and TLL1 overexpressed PC3 cells. GAPDH was used as a loading control. (**J** and **K**) Representative pictures (J) and quantification analysis (K) of migration assays in control or TLL1 overexpressed PC3 cells. Scale bar = 100 μm. \**p* < 0.05, \*\**p* < 0.01, \*\*\**p* < 0.001. Data presented in **D**, **F**, **H** and **K** are shown as mean ± SD with three times of independent assay, *p* values were assessed using two-tailed Student’s *t* tests.

**Supplementary Figure 2.**
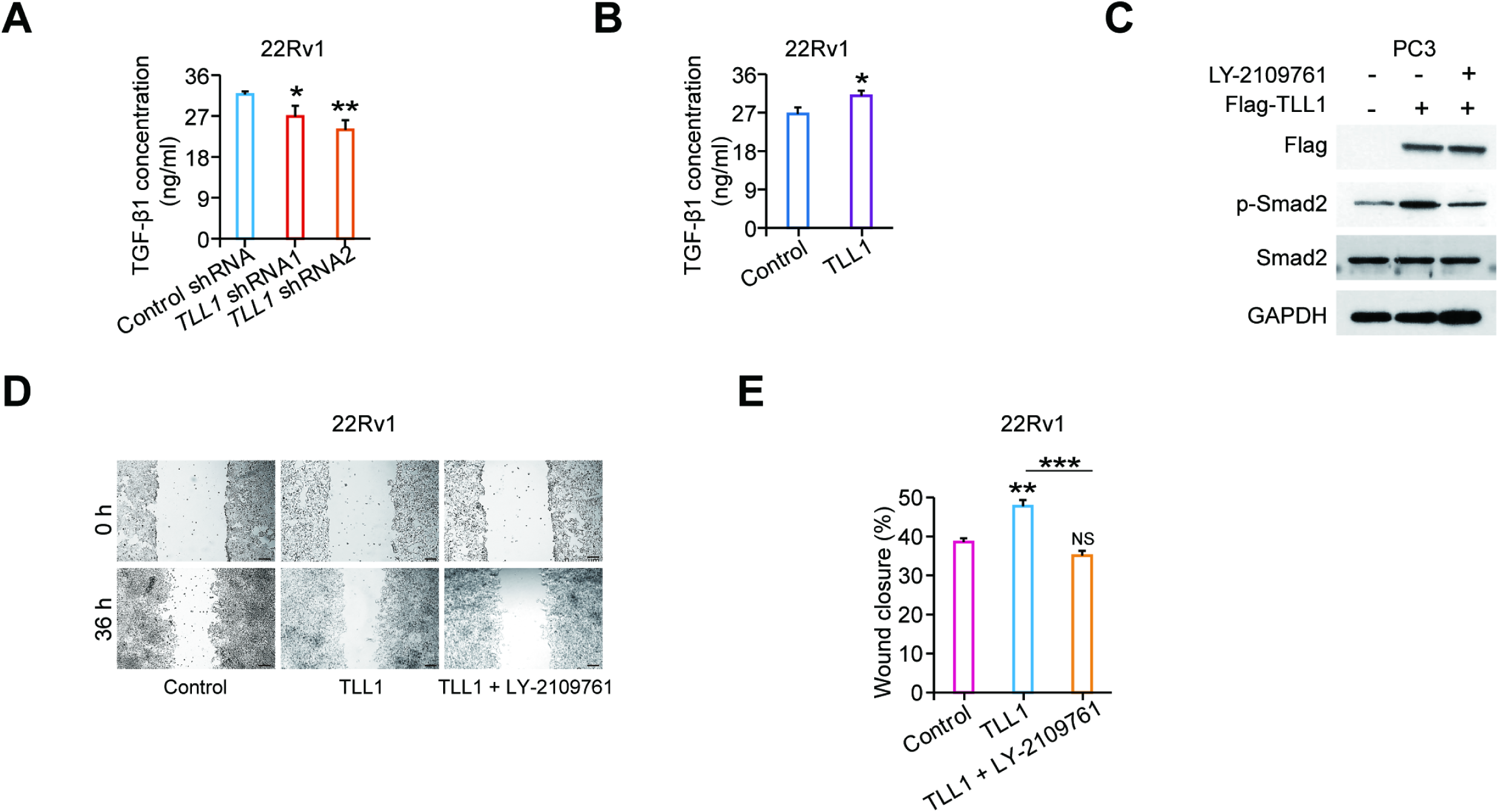
TLL1 facilitated PCa cells migration via enhancing TGF-β signaling pathway, Related to Figure 2. (A) TGF-β1 concentration was detected by ELISA assay in control and TLL1 suppressed 22Rv1 cells. (**B**) TGF-β1 concentration was detected by ELISA assay in control and TLL1 overexpressed 22Rv1 cells. (**C**) TGF-β receptor inhibitor LY-2109761(2 μM) was added in TLL1 overexpressed PC3 cells, the phosphorylated level of Smad2 were examined by western blot assay. (**D** and **E**) Representative pictures (D) and quantification analysis (E) of wound healing assays in control and TLL1 overexpressed 22Rv1 cells treated with LY-2109761 (2 μM). NS, not significant, \**p* < 0.05, \*\**p* < 0.01, \*\*\**p* < 0.001. In **A**, **B** and **E**, data shown are mean ± SD of triplicate experiments, *p* values were assessed using two-tailed Student’s *t* tests.

**Supplementary Figure 3.**
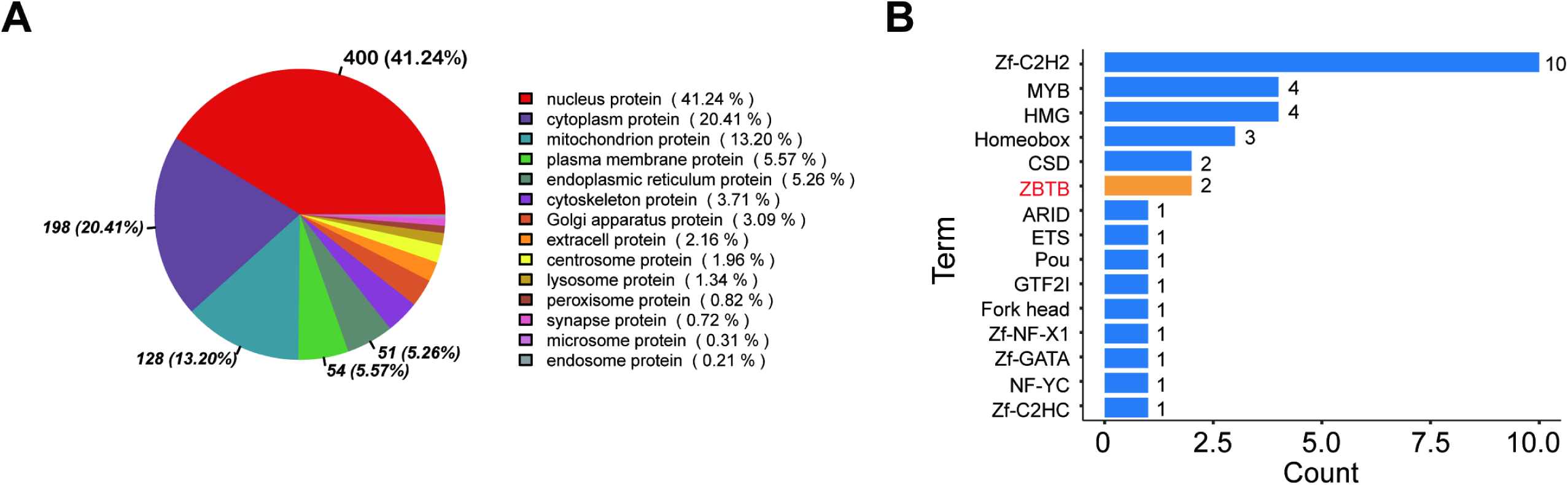
Mass spectrometry analysis identified transcription factor binding *LINC01179*, related to Figure 3. **(A)** Annotation results of subcellular localization of proteins interacting with *LINC01179*. (**B**) Transcription factor annotation results of proteins interacting with *LINC01179*.

**Supplementary Figure 4.**
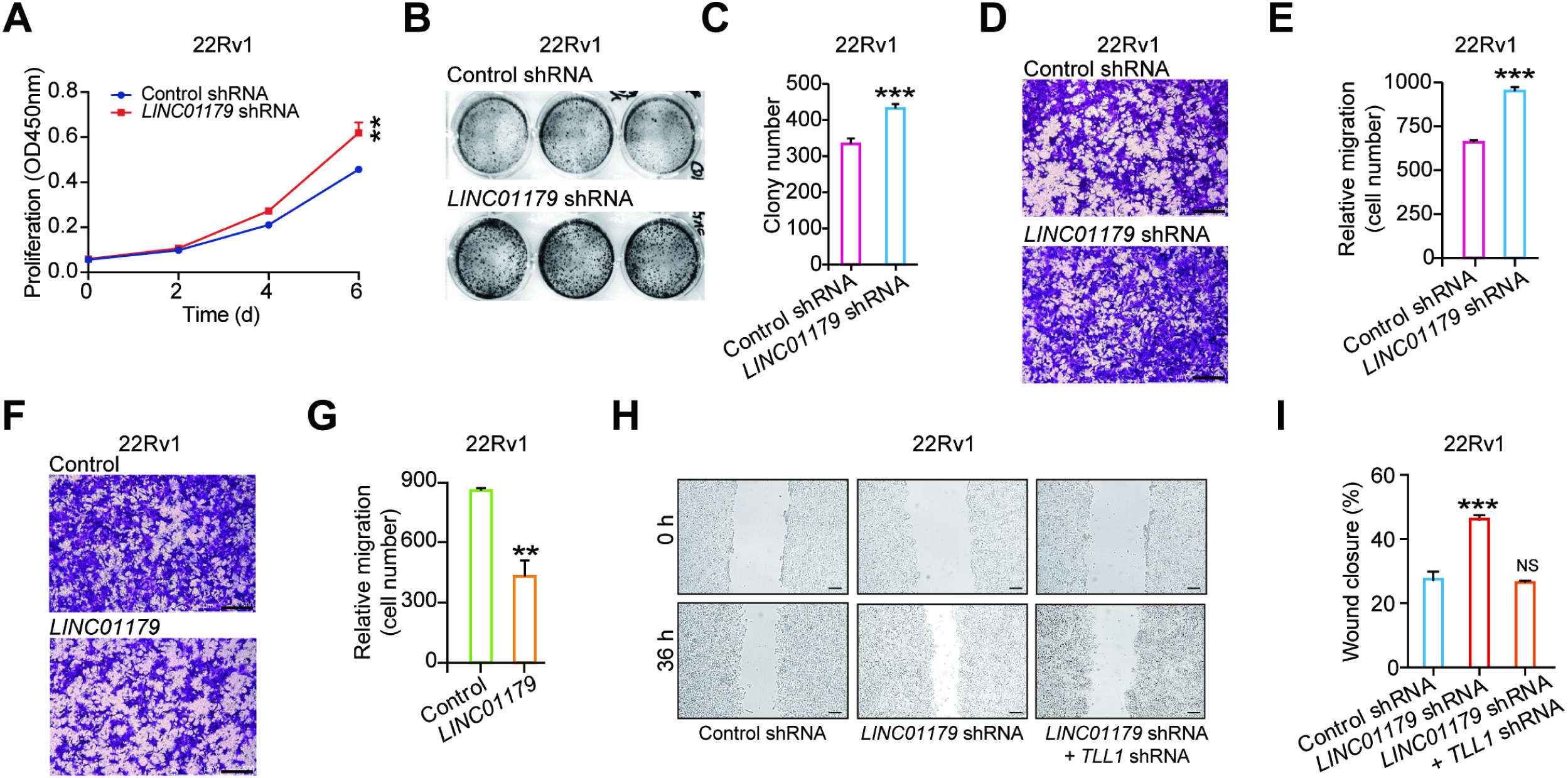
Effect of *LINC01179* on 22Rv1 cells growth and migration, related to Figure 4. (**A**) Effects of *LINC01179* depleted on cell proliferation measured by CCK8 assays in 22Rv1 cells. (**B** and **C**) Representative pictures (B) and quantification analysis (C) of colony formation assays in control and *LINC01179* knockdown 22Rv1 cells. (**D** and **E**) Representative pictures (D) and quantification analysis (E) of migration assays in control and *LINC01179* reduced 22Rv1 cells. Scale bar = 200 μm. (**F** and **G**) Representative pictures (F) and quantification analysis (G) of migration assays in *LINC01179* overexpressed 22Rv1 cells. Scale bar = 200 μm. (**H** and **I**) Representative pictures (H) and quantification analysis (I) of wound healing assays in control, *LINC01179* knockdown and *LINC01179/*TLL1 double knockdown 22Rv1 cells. Scale bar = 200 μm. Data are presented as mean ± SD of triplicate experiments. NS, not significant, \*\**p* < 0.01, \*\*\**p* < 0.001. *p* values were assessed using two-tailed Student’s *t* tests.

**Supplementary Figure 5.**
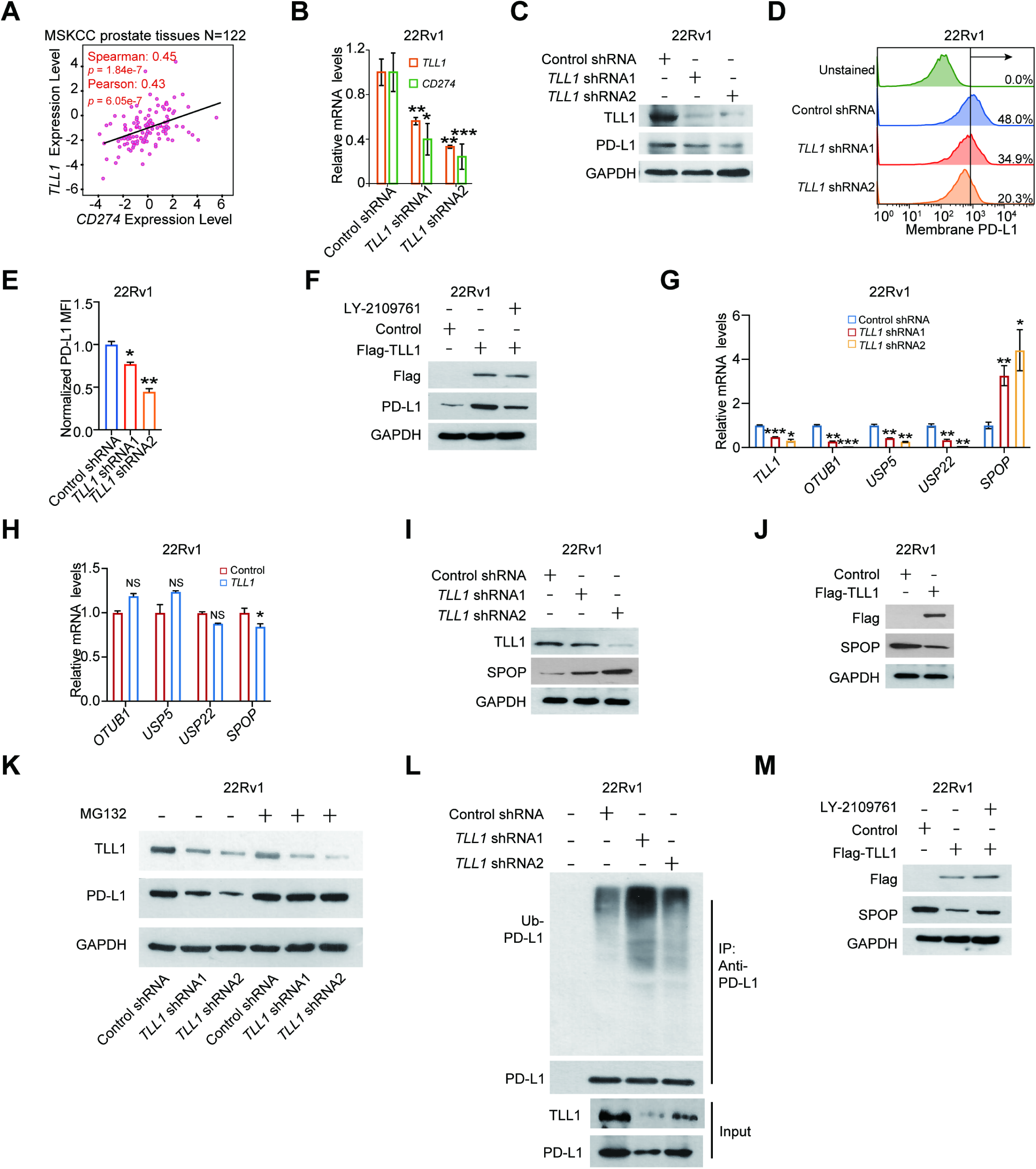
TLL1 decreased SPOP expression and increased PD-L1 expression via activating TGF-β signaling pathway, related to Figure 5. (**A**) Pearson correlation between the expression of *TLL1* and *PD-L1* in MSKCC cohort. (**B**) qRT-PCR analysis of mRNA levels of *SPOP* in control and TLL1 knockdown 22Rv1 cells. (**C**) Western blotting analysis of TLL1 and PD-L1 protein levels in control and TLL1 knockdown 22Rv1 cells. (**D**) Representative flow cytometry histograms analyzing cell surface PD-L1 in control and TLL1 knockdown 22Rv1 cells are shown. (**E**) Bar graphs showed the fold change of cell surface PD-L1 in n control and TLL1 knockdown 22Rv1 cells as measured in (**D**). (**F**) Western blotting analysis of TLL1 and PD-L1 protein levels in control and TLL1 overexpressed 22Rv1 cells treated with or without LY-2109761 (2 μM). (**G**) qRT-PCR analysis of mRNA levels of *TLL1*, *OTUB1*, *USP5*, *USP22* and *SPOP* in control and TLL1 knockdown 22Rv1 cells. (**H**) qRT-PCR analysis of mRNA levels of *OTUB1*, *USP5*, *USP22* and *SPOP* in control and TLL1 overexpressed 22Rv1 cells. **(I)** Western blotting analysis of SPOP protein levels in control and TLL1 knockdown 22Rv1 cells. **(J)** Western blotting analysis of SPOP expression levels in control and TLL1 overexpressed 22Rv1 cells. (**K**) Western blotting analysis of PD-L1 expression levels in TLL1 knockdown 22Rv1 cells or control cells treated with or without MG132 (20 μM) for 8 hours. (**L**) Control or TLL1 knockdown 22Rv1 cells were harvested for immunoprecipitation and immunoblot analysis using the indicated antibodies. Ubiquitin antibody was used to detect endogenous ubiquitinated PD-L1. Cell lysates were subjected to immunoblot with TLL1 or PD-L1 antibody. (**M**) 22Rv1 cells transfected with Flag-TLL1 or control vector were treated with or without LY-2109761 (2 μM) for 24 hours. Cell lysates were subjected to immunoblot with Flag and SPOP antibody. NS, not significant, \**p* < 0.05, \*\**p* < 0.01, \*\*\**p* < 0.001. Data are presented as mean ± SD of triplicate experiments in **B**, **E**, **G** and **H**, *p* values were assessed using two-tailed Student’s *t* tests.

**Supplementary Figure 6.**
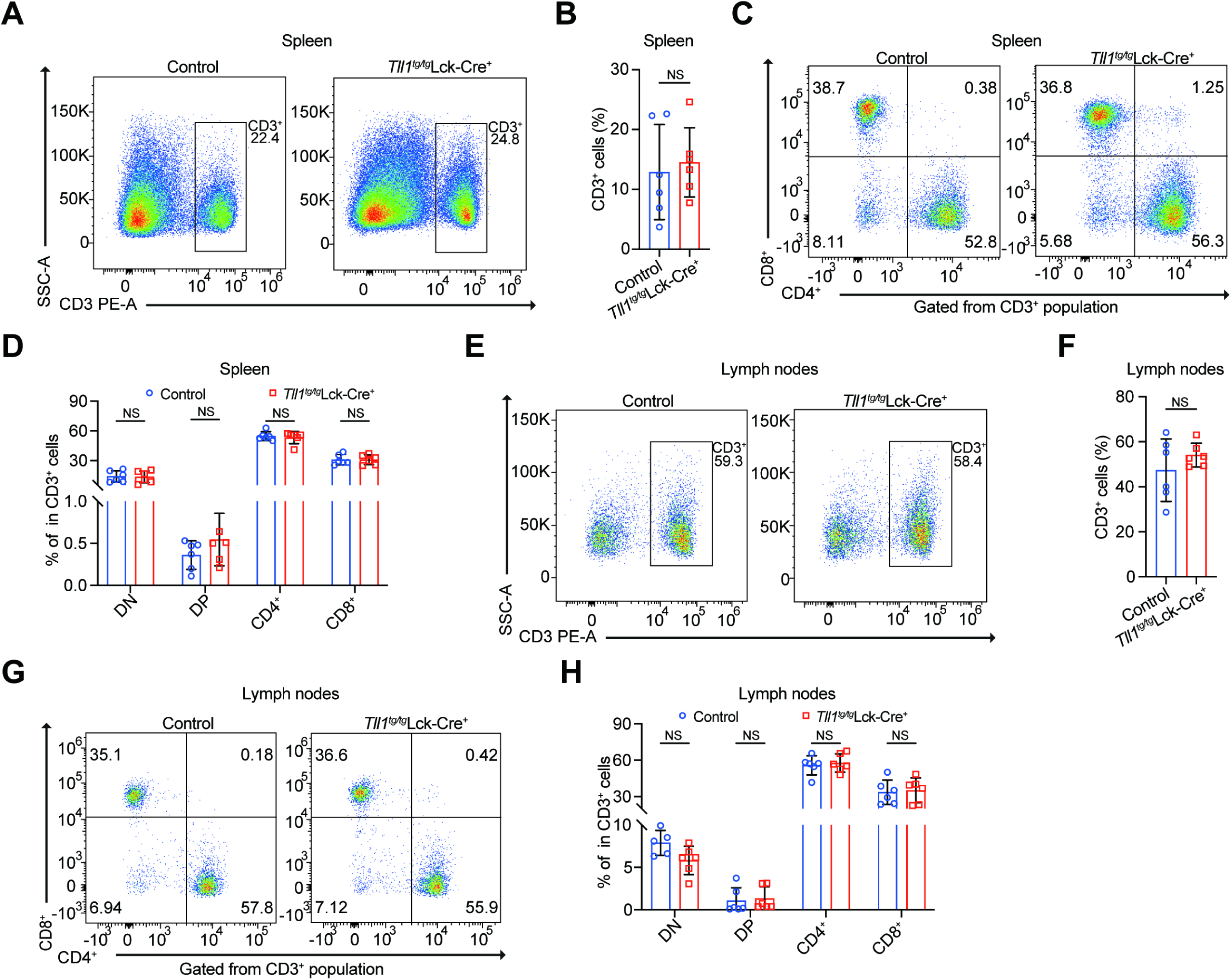
The composition of T cells in spleen and lymph nodes were normal *Tll1^tg/tg^*Lck-Cre^+^ mice, related to Figure 7. (**A**) Representative flow cytometric analysis of surface expression of CD3^+^ T cells in the spleen of control and *Tll1^tg/tg^*Lck-Cre^+^ mice. (**B**) Bar graphs of the percentages of CD3^+^ T cells in the spleen of control and *Tll1^tg/tg^*Lck-Cre^+^ mice. (**C**) Representative flow cytometric analysis of surface expression of CD3^+^CD4^+^ and CD3^+^CD8^+^ T cells in the spleen of control and *Tll1^tg/tg^*Lck-Cre^+^ mice. (**D**) Bar graphs of the percentage of CD3^+^CD4^+^ and CD3^+^CD8^+^ T cells in the spleen of control and *Tll1^tg/tg^*Lck-Cre^+^ mice. (**E**) Representative flow cytometric analysis of surface expression of CD3^+^ T cells in the lymph nodes of control and *Tll1^tg/tg^*Lck-Cre^+^ mice. (**F**) Bar graphs of the percentages of CD3^+^ T cells in the lymph nodes of control and *Tll1^tg/tg^*Lck-Cre^+^ mice. (**G**) Representative flow cytometric analysis of surface expression of CD3^+^CD4^+^ and CD3^+^CD8^+^ T cells in the lymph nodes of control and *Tll1^tg/tg^*Lck-Cre^+^ mice. (**H**) Bar graphs of the percentage of CD3^+^CD4^+^ and CD3^+^CD8^+^ T cells in the lymph nodes of control and *Tll1^tg/tg^*Lck-Cre^+^ mice. NS, not significant. In **B**, **D**, **F** and **H**, *p* values were assessed using two-tailed Student’s *t* tests.

## Notes

### Competing Interest Statement

The authors have declared no competing interest.

